# Dynamic regulation of neural variability during working memory reflects dopamine, functional integration, and decision-making

**DOI:** 10.1101/2022.05.05.490687

**Authors:** Douglas D. Garrett, Niels A. Kloosterman, Samira Epp, Vivien Chopurian, Julian Q. Kosciessa, Leonhard Waschke, Alexander Skowron, James. M. Shine, Alistair Perry, Alireza Salami, Anna Rieckmann, Goran Papenberg, Anders Wåhlin, Nina Karalija, Micael Andersson, Katrine Riklund, Martin Lövdén, Lars Bäckman, Lars Nyberg, Ulman Lindenberger

## Abstract

The regulation of moment-to-moment neural variability may permit effective responses to changing cognitive demands. However, the mechanisms that support variability regulation are unknown. In the context of working memory, we leverage the largest available PET and fMRI dataset to jointly consider three lenses through which neural variability regulation could be understood: dopamine capacity, network-level functional integration, and flexible decision processes. We show that with greater working memory load, upregulation of variability was associated with elevated dopamine capacity and heightened functional integration, effects dominantly expressed in the striato-thalamic system rather than cortex. Strikingly, behavioral modeling revealed that working memory load evoked substantial decision biases during evidence accumulation, and those who jointly expressed a more optimal decision bias and higher dopamine capacity were most likely to upregulate striato-thalamic variability under load. We argue that the ability to align striato-thalamic variability to level of demand may be a hallmark of a well-functioning brain.

---

Brain activity exhibits remarkable variability from moment to moment, exhibiting multiple dynamic signatures at every level of neural function^1,2^. Evidence is building in support of the notion that the ability to regulate neural variability provides a key signature of a well-functioning brain^2–4^. Several human studies using blood oxygen level-dependent (BOLD) signals have shown that BOLD signal variability (e.g., standard deviation, or SD_BOLD_) can be modulated within-person. On average, increases in task difficulty can drive decreases in variability^5–7^, but considerable individual differences exist. Those who can continue to elevate variability levels often exhibit a higher processing limit, maintaining better performance under higher loads, while poorer performers may hit a variability-based “cliff,” exhibiting reduced variability as load continues to increase^5,6^ (Figure 1, top). At present however, the mechanisms by which higher performing individuals are able to upregulate neural variability under load are not known.

**Figure 1:**
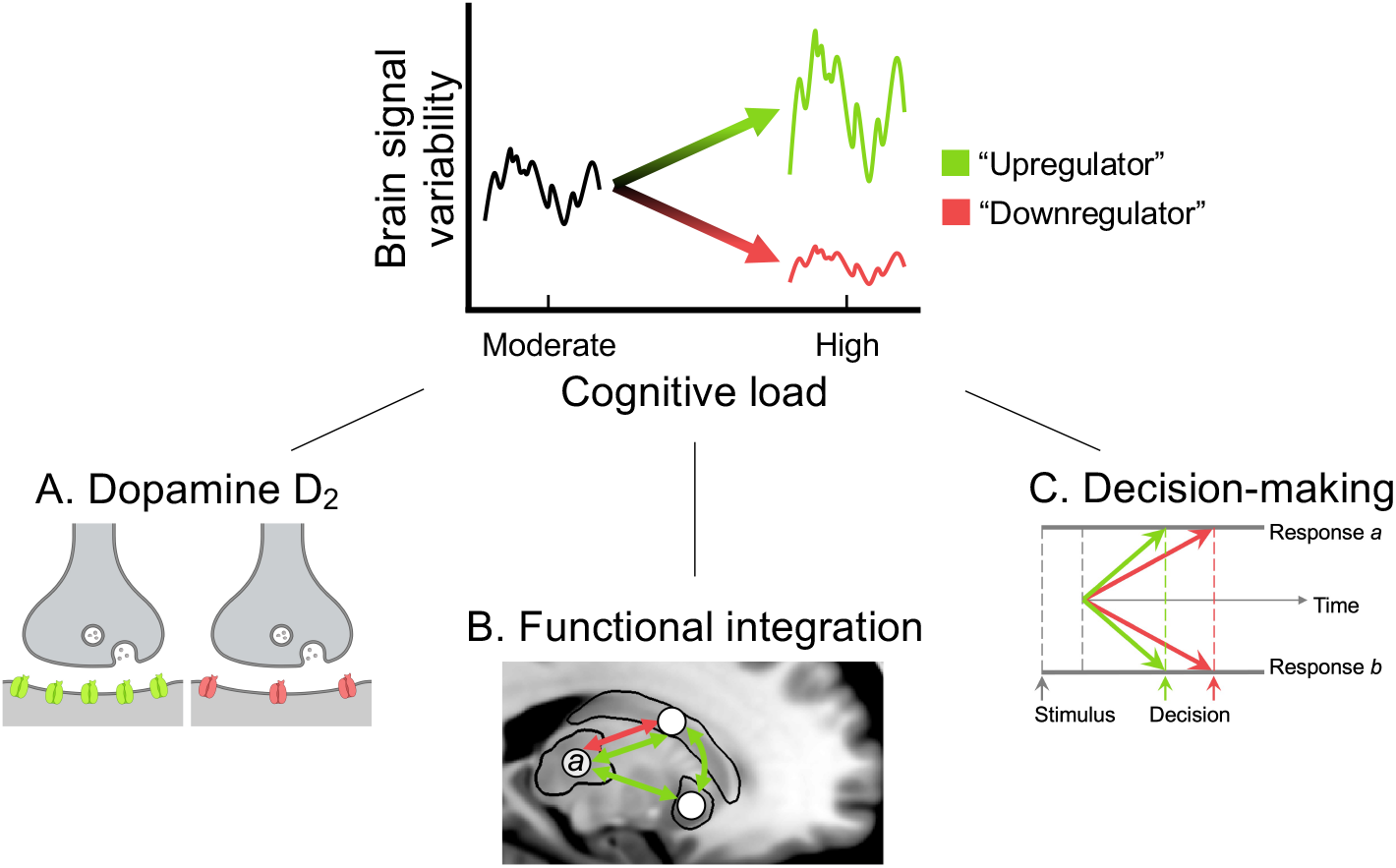
Probing individual differences in neural variability modulation with increasing cognitive load from three complementary perspectives. Older adults who can upregulate neural variability under increasing load (green) are expected to exhibit (A) greater dopamine D_2_ binding potential, (B) higher functional integration between brain regions (here, between thalamus (node a) and striatum is depicted), and (C) more effective decision-making (lower right).

One mechanism by which individuals may upregulate neural variability under cognitive load is via the dopaminergic (DA) system (Figure 1A). Although commonly associated with reward and motivation^8,9^, DA continues to gain traction as a candidate neurochemical basis for cognitively relevant aspects of brain signal variability^2^. Two human studies have shown that DA agonism can elevate the standard deviation of the BOLD signal in both younger (L-dopa^10^) and older adults (D-amphetamine^5^), while simultaneously boosting speeded responding. However, it remains unknown whether upregulation of neural variability (within-person) under greater cognitive load requires higher DA system capacity. Although the D_1_ system is inextricably linked to brain function and cognition^11–14^, D_2_ may be especially important to consider in the context of neural variability regulation under cognitive load. The D_2_ system is dominantly present in striatum and is thought to enable adaptive changes to neural processing in the face of changing environmental demands^8,13,15,16^. Crucially, striatal cells expressing D_2_ receptors project to the thalamus through the so-called “indirect” pathway^17,18^, and in turn, the thalamus provides nearly half of all excitatory input onto the striatum^19^. Among other functions, the indirect D_2_ striato-thalamic pathway allows for flexible shifting between ongoing response modes and the processing of new or salient cognitive demands^17,18^. Given that the striato-thalamic system also appears to be central for understanding how local (region level) variability reflects brain function overall^20,21^, the D_2_ system may provide an essential window into the origin and functional importance of temporal variability in the striato-thalamic system, especially as cognitive demand varies. We presume that the ability to upregulate striato-thalamic variability under load requires D_2_ capacity to meet demand.

Beyond DA, a second lens through which we can probe how individuals may upregulate brain signal variability under load is by examining “functional integration” (common temporal variation) amongst the neural populations expressing neural variability (Figure 1B). It has been proposed that dynamic neural activity at the regional level may largely reflect summed synaptic inputs^22–25^, potentially linking local dynamics to communication between regions. Indeed, more disconnected biological systems are less dynamic across moments^26^ and animal and computational work has shown that the majority of apparent “noise variation” is shared across neurons that are similarly functionally tuned^24,27^. Our recent resting-state fMRI work in humans has shown that higher temporal variability *within* a region reflects greater (i.e., lower dimensional) functional integration *between* regions across moments^20^. Critically, higher temporal variability in thalamus and striatum were among the strongest markers of functional integration across the entire brain, both cross-sectionally and longitudinally across the adult lifespan^20,21^. In simple terms, as striato-thalamic activity fluctuates during resting-state, the brain functionally integrates across moments. However, it is unknown whether local variability and functional integration are jointly regulated in the face of varying cognitive demands, and if so, whether the striato-thalamic system remains of central importance. We anticipate that individuals who can better upregulate striato-thalamic BOLD variability will also increase functional integration under load.

Third, a comprehensive understanding of the capacity to upregulate signal variability under load requires in-depth assessment of the cognitive processes giving rise to task-relevant behavioral parameters. Despite substantial progress in understanding how brain signal variability reflects traditional estimates of cognition in humans^2^ (e.g., reaction time; accuracy), it is poorly understood how neural variability (especially in the striato-thalamic system) reflects components of decision-making under varying cognitive demands. Computational models of behavior such as the drift diffusion model (DDM) conceptualize decision making as the accumulation of noisy evidence over time into an internal decision variable until a decision boundary is reached^28,29^. Such models separate non-decision (e.g., motor) from decision-related components (e.g., rate of evidence accumulation, or *drift rate*) and can also estimate the extent to which participants tend towards certain choice alternatives. Recent work suggests that those who can modulate evidence accumulation with increasing cognitive demand^30^ and adjust their decision criteria when required also express greater EEG-based variability^31^. However, spatially specific (especially striato-thalamic) signatures of how neural variability reflects evidence accumulation and decision criteria under cognitive load remain unknown. We expect that individuals who can better maintain their rate of evidence accumulation and who express optimal decision criteria in the face of rising task demands may be more likely to upregulate neural variability under load (Figure 1C).

## The current study

In the present study, we test for unique associations between brain signal variability regulation and our three research targets (D_2_ capacity, functional integration, and decision-making; Figure 1), specifically within the domain of working memory. Neural variability has proven highly responsive to working memory demands^5,7,10,32,33^, dopamine levels^5,10,12^, and the striato-thalamic system in general^5,20,21,32^. However, it is not yet known whether one’s capacity to *regulate* neural variability in the face of varying working memory load is associated with any of our research targets. In particular, computational models of decision making have not yet been deployed to understand how working memory demand drives behavior in association with variability regulation. To these ends, we directly probed load-based variability regulation by taking a fresh look at perhaps the most commonly used parametric working memory task in cognitive neuroscience, the n-back task^34^.

On this classic task, participants are asked to respond “yes” or “no” whether the current stimulus is the same as that seen *n* positions back in the set. When *n*>1, participants must not only store the immediately presented item, but also the sequence of stimuli seen^35^, creating a complex and demanding interplay between multiple cognitive processes, including attention, updating, maintenance, and inhibition^35,36^. If one can dynamically utilize these various processes across moments, temporal variation in the associated brain regions may be increased (top of Figure 1, green trace)^5,7^. However, if an individual cannot deploy such processes as capacity limits are reached (e.g., at 3-back), brain signal variability may instead compress (e.g., top of Figure 1, red), perhaps manifesting in greater behavioral losses (e.g., reduction in evidence accumulation).

To date however, precisely how the *n*-back task invokes cognitive demand is not known. Beyond the general level of working memory “demand” that n-back parametrically invokes^35^, we argue that a considerable yet rarely appreciated asymmetry exists in the demands associated with “yes” and “no” decisions on the *n*-back task. Specifically, correct “no” decisions may rely on a simple determination of stimulus set membership (“Did I see this stimulus before?”), whereas correct “yes” decisions likely require the reinstatement of details of the remembered item (“I saw it, but was it n positions back?”)^37–39^. It is plausible that those who better remember the stimulus set may more efficiently reach these simpler “No” decisions (i.e., that they are more efficient at “knowing not”^37,38^). However, as load increases, we presume that only top performers able to upregulate neural variability can maintain such an ability. We leverage the drift diffusion model to capture the effects of these various decision components on evidence accumulation in the *n*-back task^40^, in turn providing a new view of how working memory demand may drive neural variability regulation.

Using the world’s largest dataset containing both dopamine positron emission tomography (PET) and task-based functional magnetic resonance imaging (fMRI; *n* = 152 adults), we examined how our three targets (dopamine D_2_ capacity, functional integration, and drift diffusion modelling of behavior) jointly account for one’s ability to modulate fMRI-based BOLD variability under increasing working memory load. We show that greater upregulation of striato-thalamic variability is characteristic of individuals who: (i) exhibit higher D_2_ binding potential, (ii) increase functional integration, and (iii) more effectively leverage evidence accumulation processes (drift rate and drift criterion) under heightened load.

## RESULTS

### Target 1: Higher upregulation of striato-thalamic SD_BOLD_ under load uniquely reflects higher D_2_ binding potential

We first asked whether subjects who upregulate SD_BOLD_ to rising cognitive demands (i.e., from 2- to 3-back; see Methods for rationale) indeed have higher non-displaceable D_2_ binding potential (BP_ND_; assayed with ^11^C-raclopride-PET at rest). A behavioral partial least squares (PLS) model linking SD_BOLD_ modulation and D_2_ BP_ND_ revealed that individuals with higher D_2_ binding also expressed greater upregulation of SD_BOLD_ from 2- to 3-back (*r* = .30 (bootstrap 95% CI = .17, .43), *p* = 1.46e-4; see Figure 2A). This effect was most evident in dorsal and ventral striatum as well as in thalamic nuclei known to project to prefrontal cortex (medial dorsal and “motor thalamic” nuclei), as well as the intralaminar nuclei^41^ (Figure 2B). See Figure S1 for a full axial view of all nuclei.

**Figure 2:**
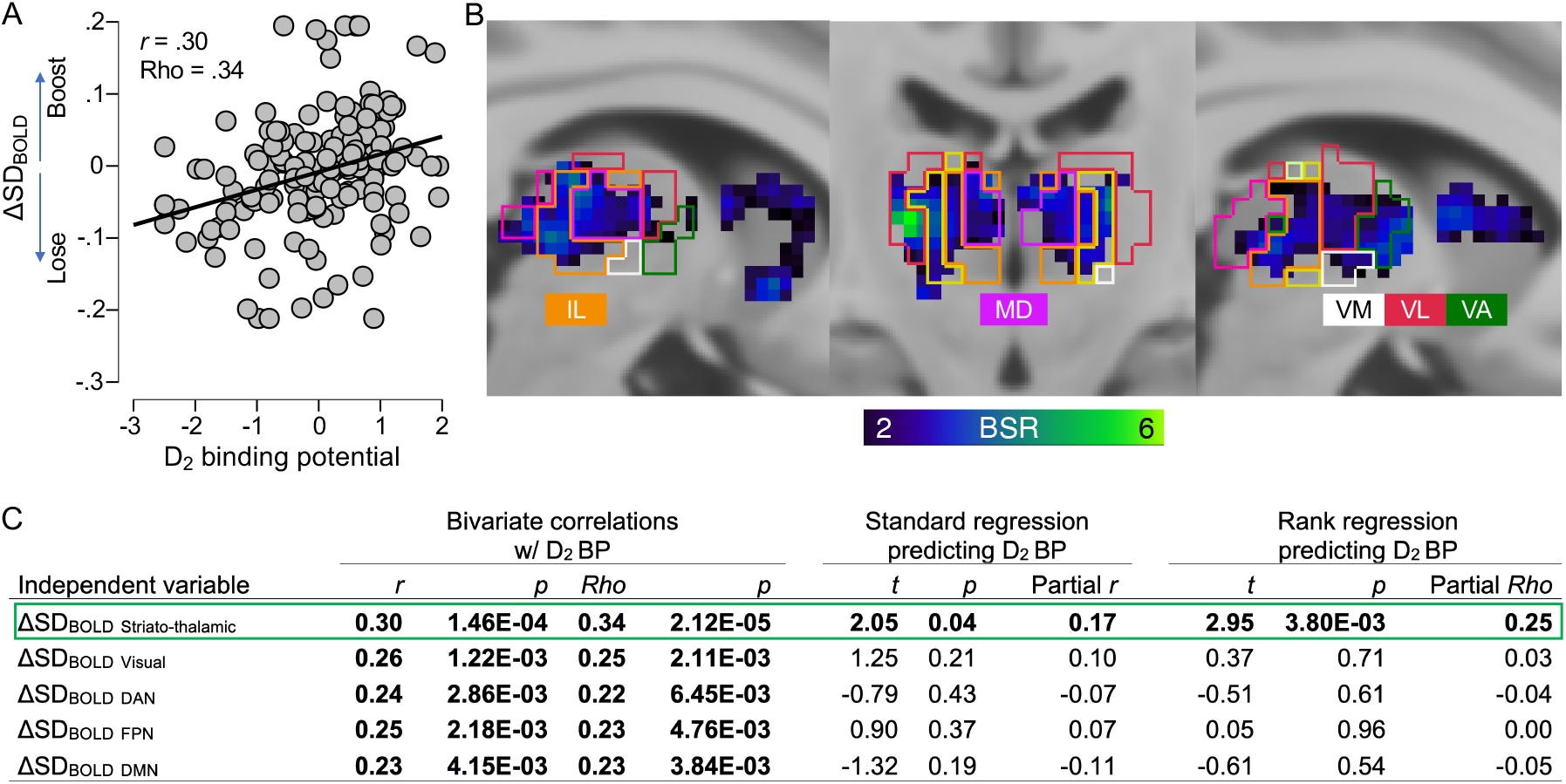
Moment-to-moment brain signal variability in the striato-thalamic system under working memory load and its association with dopamine D_2_ binding potential (BP_ND_). (A) PLS model result showing that those who upregulate SD_BOLD_ under load also have higher D_2_ BP_ND_. (B) Overlay of the Morel nucleic atlas showing key results for the intralaminar (IL), mediodorsal (MD), and “motor” (ventro-medial (VM), -lateral (VL), and -anterior (VA)) thalamic nuclei. MNI coordinates: left and middle panel = 8, -18, 4; right panel = -8, -18, 4. BSR = bootstrap ratio (higher values = more robust effects; see Methods). (C) Associations between SD_BOLD_ modulation and dopamine D_2_ binding potential. Positive bivariate effects were present in every network (see also Figure S2 for all scatter plots), but a partial effect was unique to the striato-thalamic system using both standard linear regression and rank regression (i.e., all variables present in the standard regression were rank transformed prior to model run).

We then probed whether this positive association between SD_BOLD_ modulation and D_2_ BP_ND_ was unique to the striato-thalamic network. Although SD_BOLD_ modulation was significantly bivariately correlated with striatal D_2_ BP_ND_ in every network examined (Figure 2C, left; see Methods for rationale for network choices), regression analyses revealed that this effect remained significant *only* in the striato-thalamic system when controlling for all other networks (Figure 2C, right), highlighting a distinct link between striatal D_2_ BP_ND_ and striato-thalamic SD_BOLD_ modulation over and above the rest of the brain.

### Target 2: Upregulation of SD_BOLD_ under load consistently reflects striato-thalamic functional integration

We then examined whether joint upregulation of SD_BOLD_ and functional integration (estimated via PCA dimensionality; see Methods) in the striato-thalamic system occurred with increasing cognitive load. To do so, we first correlated the DA-driven striato-thalamic ΔSD_BOLD_ brain score from the previous analysis (Figure 2) with the change in striato-thalamic PCA dimensionality from 2- to 3-back (ΔPCAdim). We found a strong negative correlation (*r* = -.71, *p* = 4.92*10^−25^; see Fig 3A), indicating that as working memory load increased, when regions within the striato-thalamic system functionally integrated in time (i.e., became lower dimensional), the time series variability of those same regions also increased. We then tested the extent to which associations between ΔSD_BOLD_ and ΔPCAdim were unique to the striato-thalamic system. First, we noted that negative associations between ΔSD_BOLD_ and ΔPCAdim were always present within each network (Figure 3B, diagonal). Strikingly however, greater load-based functional integration in the striato-thalamic network uniquely predicted upregulation of SD_BOLD_ in *every other network* (Figure 3, green box), but modulation of functional integration in non-striato-thalamic networks was *not* significantly associated with modulation of striato-thalamic SD_BOLD_ (Figure 3B, red box). These results highlight the importance of striato-thalamic functional integration for understanding working memory load-based modulation of brain signal variability across the entire brain.

**Figure 3:**
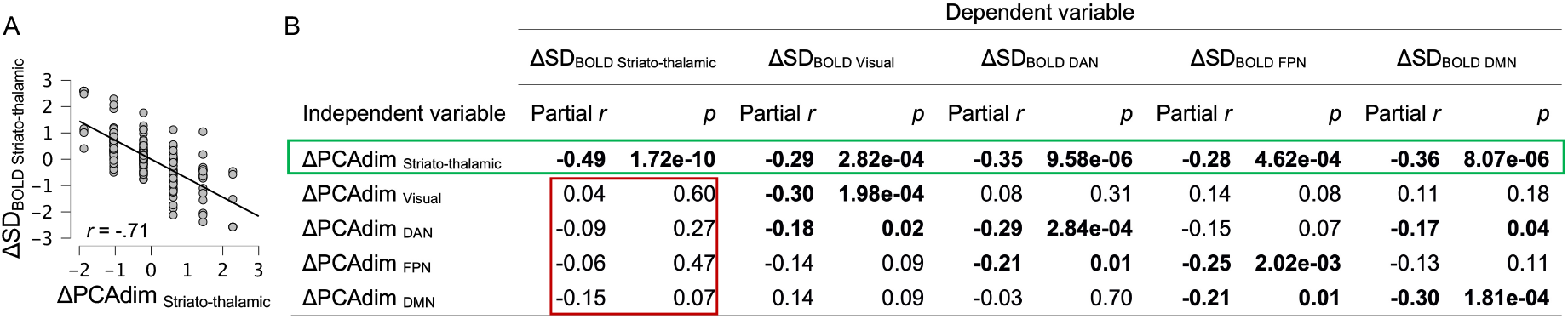
Shifts in striato-thalamic network dimensionality are uniquely associated with shifts in SD_BOLD_ in every network examined. ΔPCAdim = change in PCA dimensionality (lower values represent higher functional integration from 2- to 3-back). (A) Bivariate correlation between ΔPCAdim and ΔSD_BOLD_ in the striato-thalamic system. (B) Regressions linking ΔPCAdim to ΔSD_BOLD_ in each network. Green box: A unique effect of striato-thalamic ΔPCAdim on ΔSD_BOLD_ exists in every single network. Red box: There are no unique effects of any non-striato-thalamic ΔPCAdim measure on striato-thalamic ΔSD_BOLD_.

### Target 3a: Drift diffusion modelling of behavior

Before examining how behavior (in concert with DA and functional integration) may be associated with individual differences in SD_BOLD_ modulation under working memory load (Target 3), we first assessed the impact of rising demands on behavior using drift diffusion modelling.

#### Canonical DDM parameters

To investigate which components of the decision process were affected by the n-back manipulation, we fit the drift diffusion model (DDM) to the behavioral data^29^ using hierarchical drift diffusion modelling (HDDM)^42^. Most applications of the DDM (Figure 4A) model a binary decision (here, a “yes” or “no” decision) as the accumulation of evidence towards one of two choice boundaries. Three parameters are typically examined: the rate of sensory evidence accumulation over time (drift rate); response caution, defined as the distance between the two bounds (boundary separation), and; time spent encoding the evidence and executing the motor response (non-decision time). We found that when shifting from moderate (2-back) to high (3-back) working memory load, drift rate and boundary significantly decreased, while there was no effect for non-decision time (Figure 4B; see Table S1 for descriptives and RMANOVA results for each parameter). Also see Figure S3 (distributions) and Table S1 (descriptives and RMANOVA results) for accuracy and RT effects.

**Figure 4:**
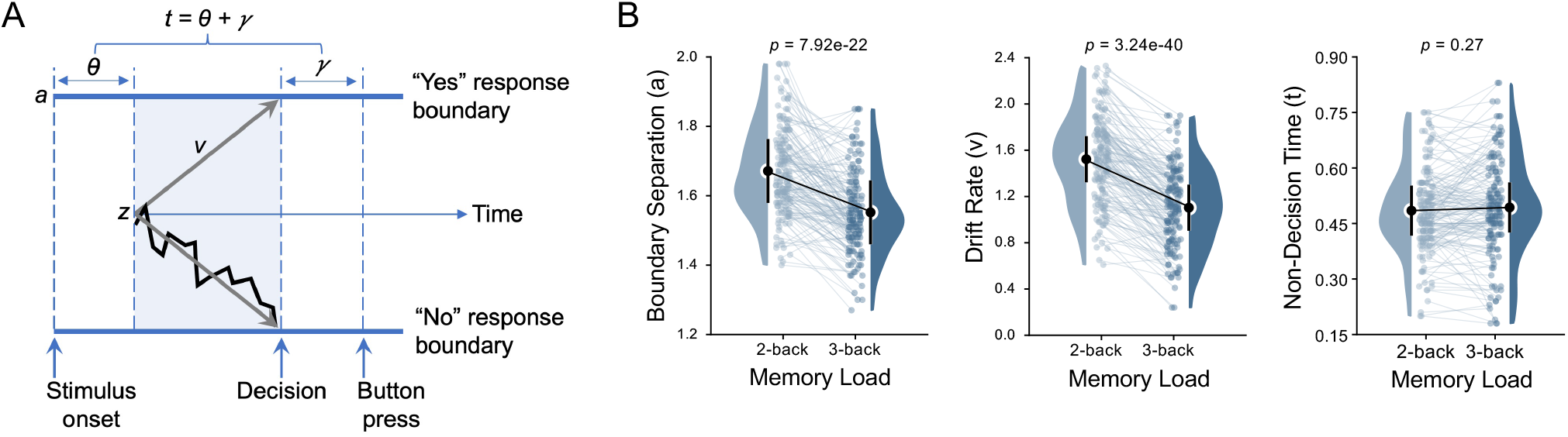
Drift diffusion model of n-back. (A) Depiction of DDM of the n-back task. a = boundary separation; v = drift rate; t = non-decision time (the sum of sensory encoding (θ) and post-decision motor components (*ϒ*)); z = starting point. (B) Canonical DDM parameter estimates by load level. Error bars represent ± 1 SD (within-subjects), calculated with the Rmisc package, using the method from Morey (2008).

#### A case for asymmetrical decision-making on the n-back task

Close consideration of the demands associated with n-back decisions reveal an asymmetry in what is required to make ‘yes’ and ‘no’ responses. Specifically, we propose that this task could be considered to have (at least) two different decision stages^39^. At stage 1, a simple determination of set membership is required (i.e., “have I seen this digit or not?”), while at stage 2, a more specific digit order retrieval process is required to compare the stimulus to that seen n-back (see Figure 5A). If the digit is new (e.g., a “6”), a “no” response can immediately be selected, whereas if the digit is recognized as part of the set already seen (e.g., a “4”), then stage 2 is invoked, requiring extra processing resources.

**Figure 5:**
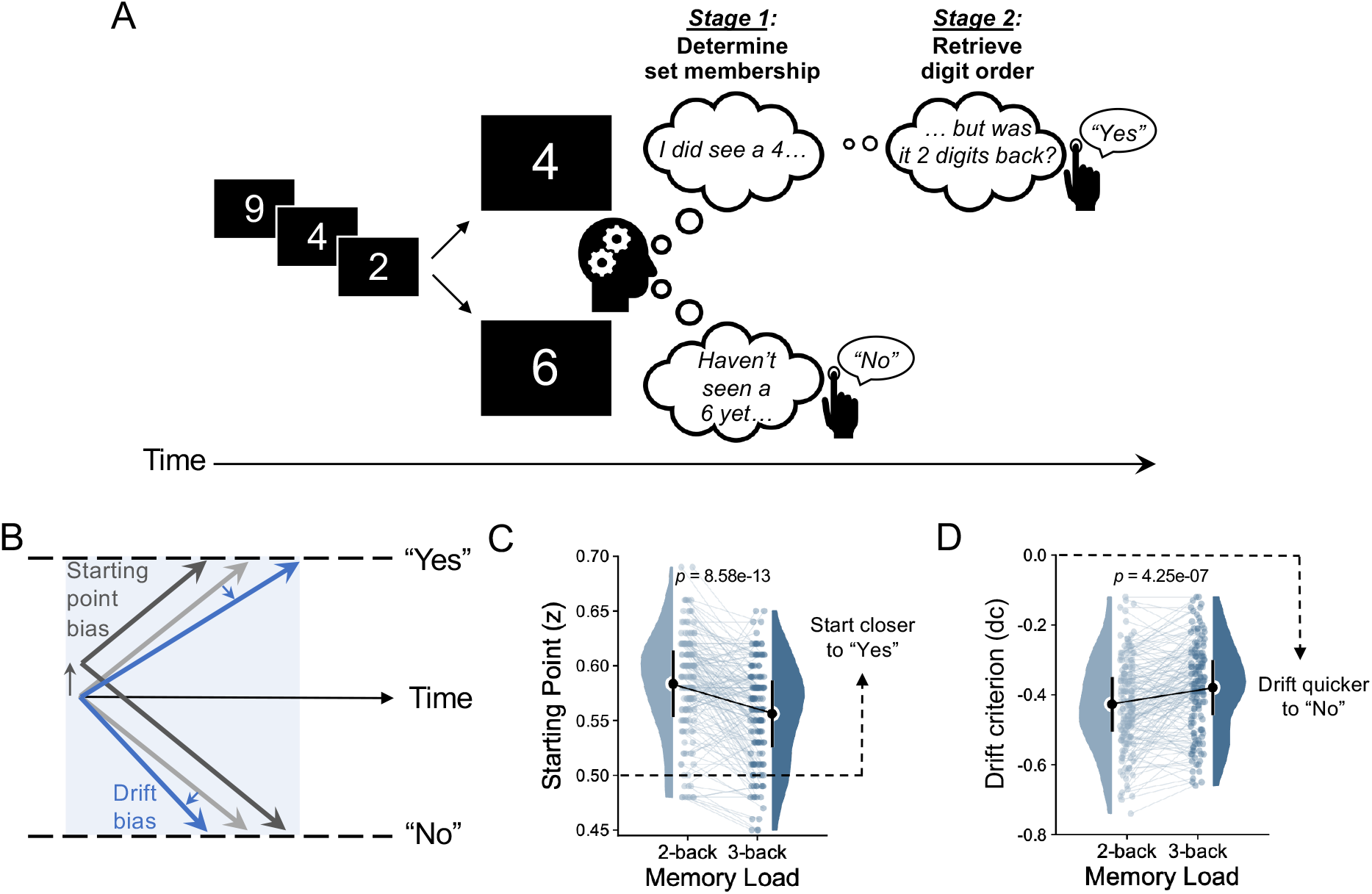
The emergence of decision bias during the n-back working memory task. The n-back task could be considered to have (at least) two different decision stages. At stage 1, one is required to determine whether an item was previously seen or not, while at stage 2, a more specific digit order retrieval process is required to compare the stimulus to that seen n-back. (A) Example in which the current stimulus is either new or previously seen relative to the set of digits already processed. If the digit is new (e.g., a “6”), a “no” response can immediately be selected. If the digit is recognized as part of the set already seen (e.g., a “4”), then stage 2 is invoked, requiring extra decision time. Decision bias in this context could present in two plausible forms. (B) Starting point bias: Knowing that “yes” responses are more resource-demanding, participants could shift their starting point towards the yes boundary to offset heightened resource demands (blue trace). Drift bias: Alternatively, a quicker/more accurate response to novel/non-target items could be captured by a quicker drift to the “no” decision boundary (blue trace). (C) Starting point bias towards “yes” responses and (D) drift bias towards “no” responses were both reduced under higher working memory load. Error bars represent ± 1 SD (within-subjects), calculated with the Rmisc package, using the method from Morey (2008).

There are (at least) two possible ways in which such asymmetrical n-back decisions may be captured by the DDM when the two decision boundaries are set to represent the actual choices (“yes” and “no”). First, by recognizing that “yes” responses require more deliberation and thus have slower evidence accumulation, participants could strategically shift their starting point of evidence accumulation towards the “yes” boundary to offset heightened memory demands. In the DDM framework, this can be estimated as a “starting point bias” that is implemented before the decision process begins (Figure 5B, dark grey trace). Second, via an alternative mechanism, higher performing participants who keep closer track of the set of within-block stimuli should be better able to recognize incoming stimuli as not being part of the set. The DDM could capture this phenomenon by estimating whether the rate of evidence accumulation is equal between “yes” and “no” responses (i.e., whether a so-called “drift criterion” exists^29^). If a stimulus is detected as new within the block, this drift criterion should lead to a more efficient decision termination at the “No” boundary (Figure 5C, blue trace). On the other hand, if the item is detected as a target (match), Stage 2 of the decision process should emerge (see Figure 5A), leading to a relatively less efficient termination at the “Yes” boundary (Figure 5B, blue trace). Crucially, unlike a starting point bias, which may be considered a general strategy (or pre-decisional bias) to offset resource demands during n-back, drift bias can *only* emerge when a stimulus has been presented, thus representing an intra-decisional form of bias directly reflecting differential evidence accumulation.

Our results indicated that a DDM including both bias terms provided a better fit to the n-back behavioral data than a standard DDM including only boundary, drift rate, and non-decision time (Deviance Information Criterion (DIC); basic model = 25809; bias model = 23641; see Figure S4 model fits for each subject for the two n-back conditions). Results indicated a robust starting point bias towards “yes” responses (Figure 5C) at 2-back (One sample t-test against zero bias (i.e., starting point bias = .50), *t* = 22.33, *p* = 1.05e-49) that tapered at 3-back (Figure 5C; *t* = 15.17, *p* = 3.72e-32). The average within-subject reduction in bias with load was also robust (*F* = 61.11, *p* = 8.58e-13, *eta*^*2*^ = .29). Next, we found a substantial drift criterion towards “no” responses in every single subject (Figure 5D) at 2-back (one sample t-test against zero bias (i.e., drift criterion = 0), *t* = 40.94, *p* = 1.21e-83) and at 3-back (*t* = 36.62, *p* = 5.36e-77). This suggests that subjects indeed leveraged the relative ease of “no” decisions on this task. The average within-subject effect was also robust (*F* = 27.97, *p* = 4.25e-7, eta^2^ = .16). Finally, in line with the idea that these two bias terms should capture distinct (pre- vs intra-decisional) aspects of n-back behavior, we noted only modest correlations between starting point bias and drift criterion at both 2-back (*r* = -.16, *p* = .06) and 3-back (*r* = .12, *p* = .16).

#### Associations between DDM parameters and offline measures

To better understand the behavioral relevance of our estimated DDM parameters, we correlated each estimated 2- and 3-back parameter with two composite measures of offline (outside of the scanner) cognitive performance, a Working Memory factor and a general Speed factor (see Methods). The two most striking sets of effects were for drift rate and drift bias (see Table S2 for all correlations). At both 2- and 3-back, higher drift rate was associated with higher offline Working Memory and faster offline Speed factor scores (2- back, *r* = .37, *p* = 2.59e-6; 3-back, *r* = .30, *p* = 1.34e-4). More negative drift bias (i.e., stronger bias towards “no” responses) was also associated with higher offline Working Memory and faster offline Speed factor scores (2-back, *r* = -.40, *p* = 2.57e-7; 3-back, *r* = -.29, *p* = 3.22e-4). Although more modest correlations between starting point bias and Working Memory (*r* = .23, *p* = 4.62e-3) and Speed (*r* = .18, *p* = .03) at 2-back were also present, such associations were not significant at 3-back. Further, both higher drift rate and more negative drift bias were associated with higher digit symbol task performance (from the Wechsler Adult Intelligence Scale) at both 2- and 3-back (abs *r* range = .18-.32). Finally, more negative drift bias was the only parameter significantly associated with more years of education (2-back, *r* = -.22, *p* = .006; 3-back, *r* = -.29, *p* = 2.83e-4). Thus, we found that decision processes reflecting evidence accumulation (drift rate and drift bias) were most convincingly associated with offline performance and educational attainment.

### Target 3b: SD_BOLD_ modulation under load as a function of DDM-based behavior (and its interactions with D_2_ BP_ND_ and functional integration)

As a final step to understanding the various bases on which neural variability modulation can occur, we ran a series of linear models linking modulation in DDM behavioral parameters, DA binding potential, and modulation of functional integration to SD_BOLD_ modulation under working memory load (see Methods). To help understand the joint role of these covariates of SD_BOLD_ modulation, interactions were of particular interest. We ran models with and without Cook’s distance outlier removal and results were highly comparable (see Table S3 for full model results).

#### Striato-thalamic system

In a single model, we first examined how dopamine binding, functional integration, and behavior may account for striato-thalamic SD_BOLD_ modulation. We found that higher D_2_ BP_ND_ and heightened functional integration were uniquely associated with upregulation of striato-thalamic SD_BOLD_ under load (see Table S3). We also noted a negative effect of drift bias, indicating that maintaining a drift bias towards “no” responses was associated with upregulation of SD_BOLD_. Notably however, we found no main effect of starting point bias in this model. Several key interactions were also present (Figure 6). First, participants with higher D_2_ BP_ND_ and greater ability to maintain a drift bias towards “no” responses were best able to upregulate SD_BOLD_ from 2- to 3- back. We also noted three interactions with modulation of functional integration, revealing that those who can jointly increase functional integration and (1) shrink their decision boundary, (2) maintain drift rate, and (3) avoid increased non-decision time under increasing working memory load were also better able to upregulate striato-thalamic SD_BOLD_. Individual differences in striato-thalamic SD_BOLD_ modulation can thus be understood as a joint reflection of D_2_ BP_ND_, network-level dimensionality, and a variety of decision processes invoked during working memory.

**Figure 6:**
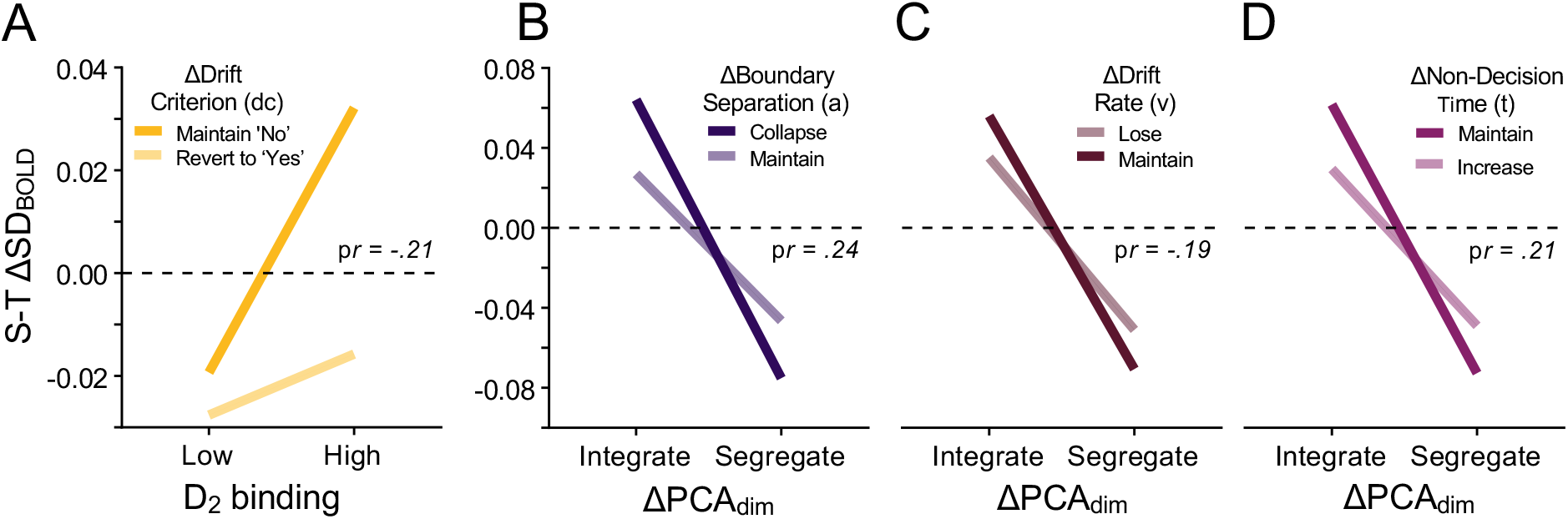
Striato-thalamic (S-T) SD_BOLD_ modulation as a function of interactions between D_2_ binding, functional integration (PCA dimensionality), and drift diffusion model parameters. Participants expressing greater upregulation of SD_BOLD_ from 2 to 3-back also (A) have higher DA and better maintain a drift bias towards “no” responses. Moreover, SD_BOLD_ upregulators also heightened functional integration while (B) collapsing their decision bounds, (C) maintaining drift rate, and (D) maintaining non-decision time under working memory load. Plots depict the visual form of two-way interactions between continuous variables, using point estimates for each variable (see Methods for details). Because point estimate error bars are not estimable from continuous variable interactions, we report the unique effect size (p*r* = partial correlation) of the interaction term estimated within each regression model; see Table S3 for all model results.

#### Beyond the striato-thalamic system

We then ran similar models (including interaction terms) for all other networks of interest (see Table S3). Most strikingly, unique main effects for *all* DDM parameters were present in a model examining the canonical working-memory-relevant fronto-parietal network (FPN; see Figure S4 for scatter plots). Those who collapsed their decision boundary, maintained drift rate, and maintained non-decision time also upregulated SD_BOLD_ under load. The FPN model was also the only one to show main effects for *both* bias terms; those with a starting point bias (z) closer to the “no” boundary and with a stronger drift bias toward “no” responses exhibited greater upregulation of SD_BOLD_ under load. However, only a single interaction was present in this model (Figure 7A); those who better upregulated functional integration and maintained drift rate also upregulated SD_BOLD_ from 2- to 3-back, just as we noted for the striato-thalamic system (Figure 6C).

**Figure 7:**
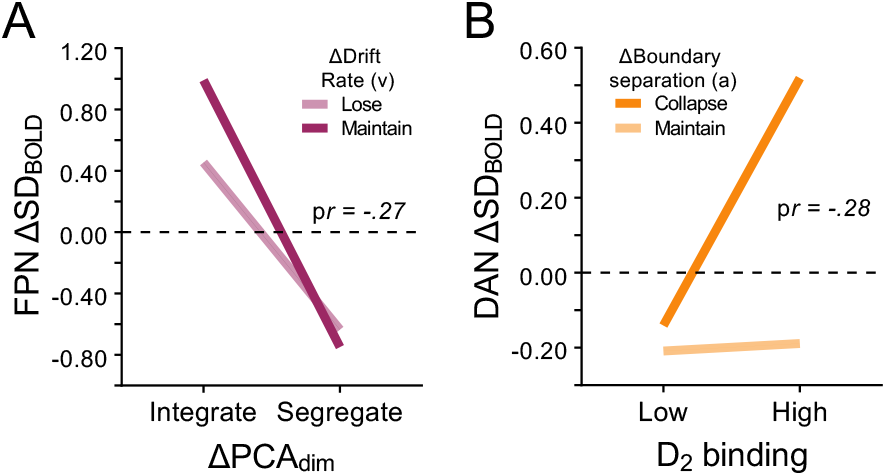
Interactions between dopamine, functional integration, and behavior beyond the striato-thalamic system. Left panel: FPN. Right panel: DAN. Plots depict the visual form of two-way interactions between continuous variables, using point estimates for each variable (see Methods for details). Because point estimate error bars are not estimable from continuous variable interactions, we instead report the unique effect size (p*r* = partial correlation) of the interaction term estimated within each regression model; see Table S3 for all model results.

For the dorsal attention network (DAN), behavioral main effects of boundary, drift rate, and drift criterion were also present (and of the same sign as for the FPN). However, only one interaction was present here too, revealing that those with higher D_2_ BP who were more likely to collapse their decision bound also upregulated SD_BOLD_ under load (Figure 7B). The visual and default mode networks showed no significant main effects of behavior or any interaction between behavior, DA and/or functional integration.

## DISCUSSION

We sought to elucidate dopaminergic, network-level, and behavioral mechanisms under which an individual can upregulate (rather than lose) brain signal variability in the face of heightened working memory demands. In many ways, the striato-thalamic system was crucial for understanding this triad of influences on signal variability modulation.

### Dopamine D_2_ binding potential is uniquely associated with modulation of striato-thalamic variability under working memory load

The positive association between D_2_ BP_ND_ and ΔSD_BOLD_ modulation was dominated by the striato-thalamic system, highlighting the centrality of this system for understanding how D_2_ is associated with moment-to-moment neural variability under heightened working-memory load. Spatially, this striato-thalamic effect was evident within a host of key thalamic regions. First, a series of thalamic regions known to project to frontal cortex were involved, including the medial dorsal (MD) and “motor thalamic” nuclei. The MD thalamus has been proposed as a key node within the generalized fronto-striato-thalamic circuitry, and as a recipient of striatal input via the pallidum, it may be critical for integrating broad-scale information within PFC during learning and memory^43–45^. Notably, MD is a key enabler of adaptive flexibility of PFC-related cognitive functions, and is broadly involved in nearly all aspects of working memory^48–52^. The “motor thalamic nuclei” (ventral medial, ventral anterior, and ventral lateral nuclei) connect directly to premotor, motor, and supplementary motor cortices in the frontal lobe^53,54^, but also to prefrontal cortex (in concert with MD nuclei) to jointly mediate associative learning, action selection, and decision-making^54^. Second, load-based modulation of SD_BOLD_ in the intralaminar (IL) nuclei was also key in our results. Previous work suggests that calbindin-positive matrix cells are prominent in the IL and other medial thalamic nuclei (e.g., ventral medial nuclei, as in the present results; see Figure 2), a cell type that projects diffusely to superficial layers across the neocortex and may constitute a thalamic “activating system” that drives effective interactions among multiple cortical areas^55,56^. The IL may also be crucial for the flexible shifting between ongoing response modes and the processing of salient cognitive demands^17,18^, here, within the context of working memory.

The notion of “flexibility” common to the actions of many of these nuclei in past work is particularly noteworthy. We found that working memory load-based modulations of brain signal variability in these thalamic nuclei were particularly sensitive to D_2_ capacity, where D_2_ is thought is also thought to enable flexible, adaptive changes to neural processing in the face of varying environmental demands^13^. Multiple forms of neural “flexibility” are theoretically required for the *n*-back task. Distinct (yet likely parallel) processes are required on a trial-by-trial basis that ensure: (1) maintenance of different memory representations over varying length delays, and (2) continuous updating of memory representations from an ongoing stream of stimulus input. Such varying demands may require “attractor plasticity,” the continual need to flexibly adjust the specific memory/representation that currently resides in attention, while still allowing established representations to exist outside of immediate attention for later recall^57,58^. Given previous conceptualizations of BOLD signal variability modulation as a general marker of system flexibility^2,3^ and the sensitivity of load-based modulation of striato-thalamic SD_BOLD_ to D_2_ capacity in the current study, future work could consider alternative working memory paradigms that allow better isolation of competing maintenance and updating stages of working memory, permitting such a “flexibility” account to be more precisely probed in relation to thalamic function.

Beyond the thalamus, dorsal and ventral striatum (bilateral putamen, caudate, and nucleus accumbens) also featured prominently in how D_2_ binding was reflected in ΔSD_BOLD_. The striatum^5,9,54–57^ has been linked to goal-directed action and motor program execution, to control of motivation, response to reward^59^ and to a variety of working memory operations^5,11,59–63^. Importantly, both dorsal and ventral striatum communicate with frontal cortex primarily via many of the same thalamic nuclei that correlate with D_2_ BP_ND_ in our results (i.e., MD, ventral lateral, and ventral anterior nuclei)^53,59^ noted above. Further, striatal D_2_ cells specifically project to thalamus through the so-called “indirect” pathway^17,18^, and in turn, the IL provides amongst the densest excitatory input onto the striatum^17,19^. Overall, our findings place the striato-thalamic system (and its frontal targets) at the core of neural substrates of how the D_2_ system may promote modulation of brain signal variability under increasing working memory load.

### As the brain to fluctuates under load, the striato-thalamic system integrates

We also found that working memory load-based upregulation of “local” signal variability was tightly coupled with increased functional integration within every network examined. This provides first human evidence that local dynamics and functional integration are closely aligned outside of the resting-state^20,21^. Crucially, striato-thalamic functional integration uniquely accounted for a sizable proportion of ΔSD_BOLD_ in every network, but functional integration in non-striato-thalamic networks was *not* independently associated with modulation of striato-thalamic SD_BOLD_. In particular, although the fronto-parietal network is broadly considered a canonical working memory network^64–67^, we found that functional integration within the striato-thalamic network was the strongest unique correlate of how moment-to-moment variation in the FPN was modulated under load. The clear presence of frontally-projecting thalamic nuclei in our results may provide a useful future basis for understanding how such an effect could be achieved in the brain. Our findings also provide a neural systems-level, working memory-based specificity to previous propositions that more disconnected, fractionated biological systems should generally be less dynamic across moments^26^. Overall, these results highlight the centrality of striato-thalamic functional integration for understanding working memory load-based modulation of neural variability across the entire brain. In short, for the brain to fluctuate under working memory load, the striato-thalamic system must be integrated (or “unified”) across moments.

### Drift diffusion modeling provides a new view of how the n-back task can be solved

Though the n-back task is among the most commonly deployed working memory tasks in cognitive neuroscience (including previous studies linking n-back to BOLD variability^5,7^), the drift diffusion model has only rarely been used to analyze n-back behavior^68,69^. In many ways, the n-back task is well suited to decouple multiple behavioral processes that may occur on this task. Our results indicated that the DDM is broadly sensitive to n-back load-based modulations at nearly all parameterized levels, offering novel insights into working memory-based decision-making. Increasing load strongly reduced the rate of evidence accumulation (drift rate), likely indexing the heightened difficulty of evidence accumulation during working memory search as demands increase and capacity limits are reached. Boundary separation also shrunk under load, suggesting a general reduction in response caution. This effect may indicate that participants recognize that 3-back is appreciably more difficult, in turn reducing their internal criterion for how much evidence is required prior to making a decision; in this way, decisions can still be made within the available response window despite increased task difficulty.

Perhaps most strikingly, the estimation of bias parameters served to capture previously unappreciated individual differences in n-back performance. Drift criterion was the most robust form of response bias in the present data; every single subject expressed a drift criterion towards “no” responses (i.e., where “no” means that the current stimulus does not match that seen n-back). Why might subjects drift more quickly toward “no” on this task? Like many previous studies on the *n*-back task, the lure rate in the current task is negligible, to prevent confusion between target and non-target items (see Methods). In this light, our drift criterion effect then becomes clear once a clear decision asymmetry inherent in the construction of the n-back task is acknowledged (see Figure 4). We have framed the n-back task as requiring two key decision stages. At Stage 1, simple set membership of the stimulus is judged. Given a low lure rate, if the subject doesn’t recognize the stimulus at any level, then “no” can be immediately selected as the item is almost certainly new. However, if the stimulus is recognized, then Stage 2 is required to specify whether the recognized stimulus was seen n positions back, requiring a more specific and resource intensive form of memory reinstatement.

We postulated that better performers in general may more efficiently reach these simpler “No” decisions (i.e., that they are more efficient at “knowing not”^37,38^). In accord with this prediction, we showed that those exhibiting a greater drift criterion towards “no” responses were more successful on a series of offline working memory and speed tasks, had higher general intelligence, and greater years of education. The ability to leverage such a drift criterion under load may thus characterize a well-performing system in general, and our findings open a new window into how cognitive “demand” on the classic n-back task may manifest.

### The regulation of neural variability is jointly accounted for by behavior, dopamine capacity, and functional integration

Beyond average DDM-based behavioral effects on the *n*-back task, our core interest here was examining how load-based modulations in SD_BOLD_ may be accounted for *jointly* by performance, D_2_ binding, and functional integration. We noted that the striato-thalamic system was again uniquely sensitive, revealing a series of notable interactions. First, under increasing working memory load, those who increased functional integration and: (a) collapsed their decision bound, (b) maintained (rather than lost) drift rate, and (c) maintained (rather than increased) non-decision time were best able to upregulate striato-thalamic SD_BOLD_ under load. These results suggest that the ability to avoid falling off a variability-based “cliff” under load may require that better performers also maintain a low dimensional, temporally “unified” striato-thalamic system.

Strikingly, the striato-thalamic system was the only network to reveal that higher D_2_ binding and better maintenance of a negative drift criterion under load combined to account for upregulation of SD_BOLD_. How might this interaction between D_2_ capacity and drift bias work? As noted above, the better that subjects keep track of the stimuli they have seen, the effectively they should recognize that a stimulus has not yet been seen. As such, previously unseen stimuli could represent a particularly *salient* (“pop-out”) signal for a well-functioning working memory system, creating the context for a more efficient drift to “no” responses to exist. Dedicated circuitry in the striato-thalamic system may permit this to occur (see Figure 8). In general, the indirect (D_2_ -based) striato-thalamic pathway allows the thalamus to interrupt ongoing cortical response modes to flexibly re-orient the brain toward particularly salient stimuli^17,18^. As summarized by Gerfen et al.^17^ and Ding et al.^18^, the intralaminar nuclei of the thalamus in particular can rapidly respond to salient stimuli, creating an immediate burst/pause function in cholinergic interneurons that suppresses the ongoing cortical drive of striatal circuits, in turn strongly biasing the striatal network towards a more flexible D_2_ regime. Higher D_2_ binding potential could then be suggestive of a higher-capacity indirect pathway system, such that the intralaminar nuclei have more options available to them when the time comes to shift from the ongoing processing of already seen items toward the processing of new (salient) stimuli. Animal work suggests that this primary thalamo-striatal pathway is also crucial for reducing interference between new and existing learning^70^ and becomes degraded in aging^71^. Such interference reduction may also be critically important for simultaneously performing the maintenance and updating components of the n-back task. Further, D_2_ striatal neurons are also remarkably excitable over a range of inputs^17,72^, producing relatively higher moment-to-moment dynamics (than D_1_) as a result. We may have captured some aspects of those dynamics via working memory load-related shifts in SD_BOLD_ in the current study.

**Figure 8:**
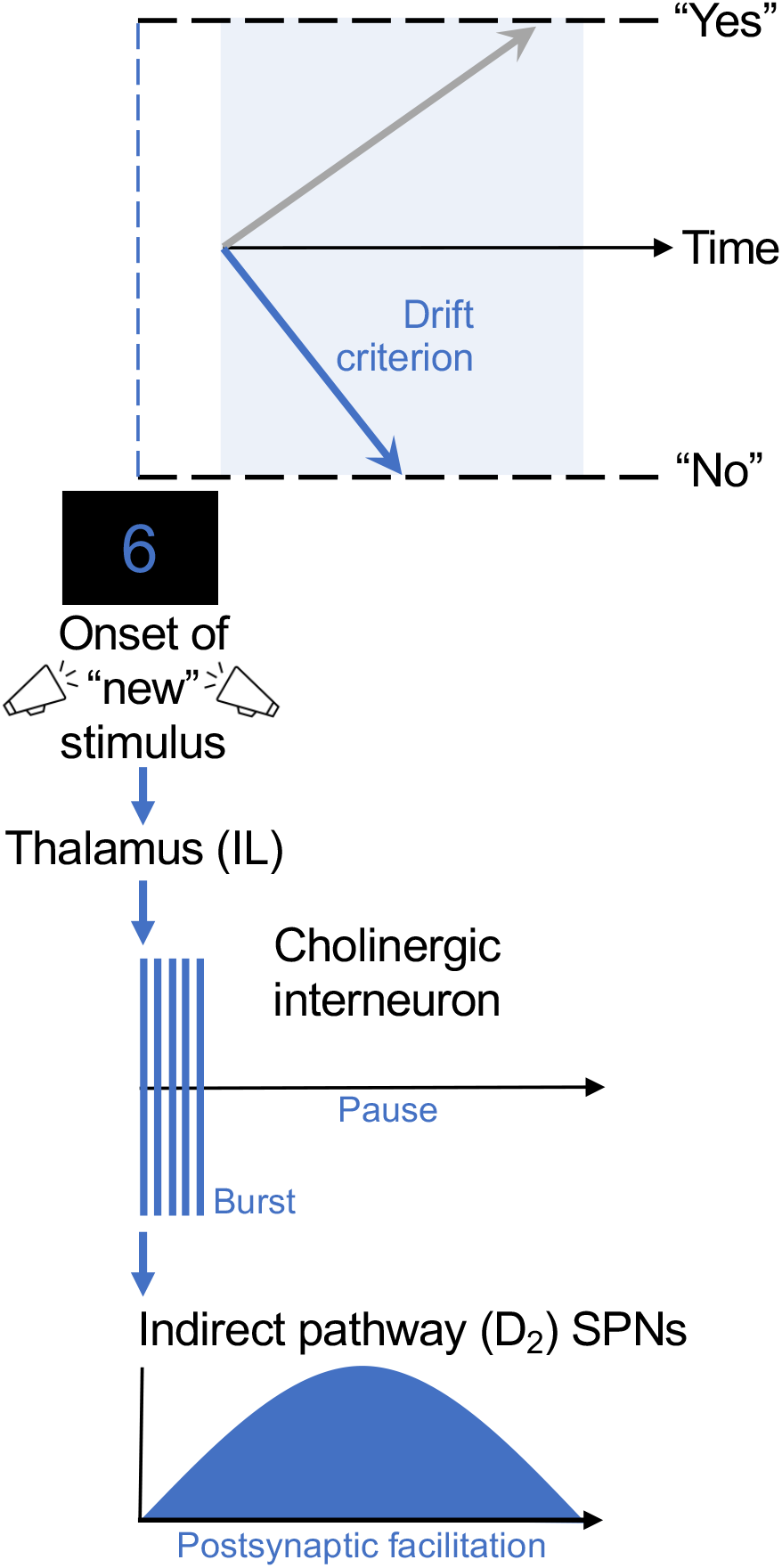
Depiction of how effective processing of new items in n-back could permit a negative drift criterion by engaging a D_2_-specific thalamo-striatal circuit. For example, at 3-back, upon onset of an unseen stimulus (e.g., a 6, after previously seeing 4-7-9-4), drift criterion towards a “no” decision (i.e., the item does not match that seen n-back) emerges. This novel stimulus acts as rapid input to the intralaminar nucleus of the thalamus, which creates a burst of activity in cholinergic interneurons, ultimately biasing spiny projection neurons (SPNs) in the striatum toward a D_2_ regime for ∼1 sec. Figure adapted from ^*17,18*^.

Interestingly, although the striato-thalamic system revealed the only robust D_2_ by drift bias interaction on SD_BOLD_, drift bias modulation was the single most consistent main effect across all networks showing brain-behavior associations (i.e., striato-thalamic, DAN, and FPN). It is notable that SD_BOLD_ modulation in frontally-projecting thalamus and the FPN and DAN networks (both with strong lateral frontal regional representation) were all sensitive to drift bias on n-back. Indeed, the FPN is perhaps the most common *a priori* target in the working memory literature and is thought generally responsible for top-down control of working memory representations^73,74^. The DAN (often termed the “task positive network”) typically becomes more active during a host of different tasks requiring externally-oriented attention. Combined, the broader fronto-striato-thalamic system overall may be a primary target for understanding how working memory-based drift bias can be implemented by the human brain. Further, SD_BOLD_ modulation in the FPN was remarkably sensitive to main effects of *all* estimated DDM parameters, supporting its general role in how working memory load-based modulation of behavior is associated with BOLD variability. Further, SD_BOLD_ modulation in the FPN was sensitive to starting point bias *and* drift criterion modulation, with the effect of each pointing in the same direction – those who start closer and drift quicker to “no” expressed greater upregulation of SD_BOLD_ under higher working memory load. We argue that such biases should serve as a key future target for undertstanding the neural dynamics of working memory.

### Future directions

There are several future directions that could further clarify how dopamine, functional integration, and behavior are associated with moment-to-moment neural variability. First, to fully interrogate how decision bias manifests in n-back/working memory, future task designs could directly manipulate the capacity for bias to emerge by manipulating the lure rate at each load level. Second, our measure of striatal D_2_ is an estimate of binding potential, gauging the capacity of the system to utilize a D_2_ regime. An estimate of real-time DA dynamics would be ideal for testing whether the indirect (D_2_) striato-thalamic pathway is preferentially engaged (e.g., over the direct (D_1_) pathway) as drift criterion is implemented under increasing working memory load. Although sub-second estimates of DA fluctuations have been achieved in Parkinson’s patients undergoing deep brain stimulation^75^, this has not yet proven possible in healthy adults. Third, alternative neuroimaging modalities, data processing routines, and variability estimation methods can sometimes reveal spatially and statistically differential results (see^2,12,33,76,77^). In particular, choice of artifact removal and normalization technique prior to estimating variability can dramatically shift results. As such, future studies should be careful to document precisely how variability is computed and interpret their results in that light. Finally, the present multi-modal results were obtained with an unusually large sample of older adults. It remains to be seen whether our effects are equally strong during earlier periods of the human lifespan. To achieve success in typical young adult samples, higher n-back load levels may be required to invoke individual differences in modulation on the level of those seen in the present study.

## Conclusions

Our work suggests that an individual’s ability to upregulate moment-to-moment variability in brain activity under working memory load is jointly associated with higher D_2_, higher functional integration, and more optimal decision-making. In particular, the striato-thalamic system appears to represent a rich nexus for understanding how these three signatures interact to determine neural variability regulation. We argue that the ability to upregulate brain signal variability under working memory load may be a crucial hallmark of an effective adult brain.

## METHODS

A comprehensive description of the recruitment procedure, imaging protocols, and cognitive and life-style assessments in the large-scale Cognition, Brain, and Aging (COBRA) study have been published elsewhere^62,63,78,79^. Here, we describe the methods directly relevant to the present study.

### Participants

The current sample consisted of 152 older adults (64-68 years; mean = 66.2, SD = 1.2; 81 women) randomly selected from the population register of Umeå, in northern Sweden. As noted in previous work on this sample^78^, exclusion criteria included suspected brain pathology, diabetes, treatment for cancer, neurological and psychiatric disorders, impaired cognitive functioning (Mini Mental State Examination < 27), and conditions that could bias the brain measurements (e.g., severe trauma, tumors), cognitive performance (e.g., severely reduced vision), or preclude imaging (e.g., metal implants). 28% of the sample was working, 18% used nicotine, and 33% took blood-pressure medications. Mean education was 13.3 years (SD = 3.5), body-mass index (BMI) was 26.1 (SD = 3.5), systolic blood pressure was 142 (SD = 17), and diastolic blood pressure was 85 (SD = 10). The sample is representative of the healthy target population in Umeå, Sweden.

### Image Acquisition

Magnetic resonance (MR) imaging was performed with a 3 Tesla Discovery MR 750 scanner (General Electric, WI, US), equipped with a 32-channel phased-array head coil. A 3D fast-spoiled echo sequence was used for acquiring anatomical T1-weighted images, collected as 176 slices with a thickness of 1 mm. TR = 8.2 ms, flip angle = 12 degrees, and field of view = 25 × 25 cm. BOLD-contrast sensitive scans were acquired using a T2*-weighted single-shot gradient echoplanar-imaging sequence. Parameters were: 37 transaxial slices, 3.4 mm thickness, 0.5 mm spacing, TE/TR = 30/2000 ms, 80 degrees flip angle, 25 × 25 cm field of view, and a 96 × 96 acquisition matrix (Y direction phase encoding). At the start, 10 dummy scans were collected. Functional data were acquired during a resting-state condition (6 min) followed by the numerical n-back WM task described above.

Positron Emission Tomography (PET) was performed with a Discovery 690 PET/CT scanner (General Electric, WI, US) during resting-state conditions, following an intravenous bolus injection of 250 MBq [11C] raclopride. Preceding the injection, a 5 min low-dose helical CT scan (20 mA, 120 kV, 0.8 s/revolution) was acquired, for attenuation correction. Following the bolus injection, a 55-min 18-frame dynamic scan was acquired (9×120s, 3×180s, 3×260s, 3×300s). Attenuation, scatter, and decay-corrected images (47 slices, field of view = 25 cm, 256 × 256-pixel transaxial images, voxel size = 0.977 × 0.977 × 3.27 mm) were reconstructed with the iterative resolution-recovery VUE Point HD-SharpIR algorithm, using 6 iterations, 24 subsets, and 3.0 mm post filtering, yielding full width at half maximum (FWHM) of 3.2 mm^80^. Head movements during the imaging session were minimized with an individually fitted thermoplastic mask attached to the bed surface. For 82% of the individuals, PET was carried out 2 days after the MR scan (average time difference between MRI and PET: 3 ± 6 days).

### In-scanner task

During fMRI scanning, participants performed a numerical n-back task. In this task, a sequence of single digits appeared on the screen. Each digit was shown for 1.5 s (within which a response must have been made), with an ISI of 0.5 s. During every item presentation, participants reported if the number currently seen on the screen was the same as that shown 1, 2, or 3 digits back. A heading that preceded each blocked condition indicated the load level. Participants responded by pressing one of two adjacent buttons with the index or middle finger to reply ‘yes, it is the same number’ or ‘no, it is not the same number’, respectively. A single fMRI run with 9 blocks for each condition (1-, 2-, and 3-back) was performed in random order (inter-block interval: 22 s). Each block consisted of 10 trials that included 4 matches (requiring a “yes” response) and 6 non-matches (requiring a “no” response). Within each block of 10 trials, the number of valid trials depended on n-back level (1-back: first trial dropped, 9 valid trials, 4 yes/5 no trials; 2-back: first two trials dropped, 8 valid trials, 4 yes/4 no trials; 3-back: first three trials dropped, 7 valid trials, 4 yes/3 no trials), yielding a total of 81, 72, and 63 valid trials per subject for 1-, 2-, and 3-back respectively. The specific stimulus/trial sequence was the same for all participants, with only two lures total (a single 2-back lure within two of the 3-back blocks). The n-back condition blocks were counterbalanced.

For the purposes of the current study, we focus primarily on the 2- and 3-back conditions. Although typically examined when the n-back task is reported in the literature, 1-back demands can easily be considered qualitatively different from 2 or 3-back. For 1-back, all that is required is for a single digit to be maintained in memory for 0.5 sec; each digit seen is simply compared to the one that comes immediately afterward. From the perspective of component processes of working memory^36^, this simple process of maintaining a single stimulus in mind has even been framed as a “focus of attention”^35,81^. For 2- and 3-back however, one must maintain a given stimulus and its temporal order in memory for multiple seconds *in the face of* ongoing updates to the string/list of numbers to be remembered. Further, performance is typically extremely high at 1-back^5,7,35,66^ and shows minimal associations with D_2_ BP_ND_ in past work^63^. With all these issues combined, we thus focused on the 2- and 3-back conditions throughout the current study.

### Behavior

#### Drift diffusion modeling of choice behavior

We fitted a drift diffusion model (DDM) to the accuracy and RT data of the 2- and 3-back conditions to quantify the dynamics of the cognitive processes underlying working memory-based decisions^29^. The DDM is a sequential sampling model that provides an algorithmic account of how the accumulation of evidence over time contributes to a binary decision process. To this end, the DDM decomposes the decision process into three basic parameters: drift rate, capturing the degree of evidence accumulation; separation of the decision boundaries representing each alternative, and; non-decision time spent on sensory encoding and motor response. The DDM has successfully been applied to the n-back task in previous work^82^. In addition to these standard parameters, we estimated two parameters related to decision bias to account for systematic preferences for either ‘yes’ or ‘no’ choices^83^, one invoked prior to the onset of the evidence accumulation process (starting point bias), and another representing a bias in the evidence accumulation process itself (drift bias). See Results for our detailed rationale for including these bias parameters in the context of n-back.

We used hierarchical drift diffusion modeling as implemented in the HDDM toolbox^42^ to estimate model parameters. To enable estimation of the response bias parameters, we fit the model using separate correct and error RT distributions for target-present and target-absent trials. This procedure is termed ‘stimulus coding’ in the HDDM toolbox, as opposed to the more common ‘accuracy coding’, where RT distributions are fit separately for correct and error trials, independent of the stimulus. The hierarchical Bayesian parameter estimation of the DDM in the HDDM-toolbox constrains single-participant parameters estimates by the group and therefore results in stable estimates also when within-subject data are limited but the number of subjects is relatively large, as in our dataset.

We ran 5 separate Markov chains for the Bayesian estimation with 5000 samples each. The first 2500 samples were discarded as burn-in, i.e., to let the sampler identify the region of best fitting values in the parameter space. The remaining 2500 samples per chain were concatenated across chains. Individual parameter estimates were then estimated from the posterior distributions. All group-level chains were visually inspected to ensure convergence. Additionally, we computed the Gelman-Rubin R-hat statistic (which compares within-chain and between-chain variance) and checked that all group-level parameters were below 1.1^84^. We also performed posterior predictive checks to assess whether the model was able to capture choice and reaction-time patterns (Fig. S4).

#### Offline measures

To better characterize the relevance of DDM parameters, we also utilized several offline measures of interest. We examined unit weighted composite scores of Working Memory (average score across letter updating, columnized numerical 3-back, and spatial updating tasks) and Speed (average across letter, number, and figure comparison) used in previous work (see ^78,85^). We also used the digit symbol subtest from the Wechsler Adult Intelligence Scale, a task capturing broad-scale aspects of processing speed, attention, associative learning, and executive function^86^.

### Image Processing

#### PET

The following preprocessing steps were performed for each subject in SPM8. The 18 frame PET scans were coregistered to the T1-image using the time-frame-mean of the PET images as source. They were then normalized to MNI-space with the subject-specific flow fields (obtained with DARTEL) and then affine transformed and smoothed via a Gaussian filter of 8mm. Normalization parameters were selected so that concentrations in the images were preserved. For determination of D_2_ BP, time-activity curves for each voxel were entered into a Logan analysis (Time frames 10-18 (18-55 minutes))^87,88^, using time-activity curves in the grey-matter parts of cerebellum as reference. Regions of interest were delineated with the FreeSurfer 5.3 segmentation software ^89–91^. Median BP to non-displaceable tissue uptake (BP_ND_) data were extracted for all regions of interest based on the subcortical parcellations in Freesurfer and the Desikan-Killiany atlas ^92^ for extrastriatal regions.

Our specific estimate of striatal D_2_ BP_ND_ was taken from previous work on the COBRA dataset using structural equation modeling (SEM) to model between-person differences in D_2_ availability in a variety of striatal and extrastriatal regions^93^. Briefly, [^11^C]raclopride BP_ND_ of the left and right caudate and putamen was estimated as a single latent striatal factor. Single subject values on this factor were calculated using regression-based estimation of factor scores; see Papenberg et al.^93^ for further details.

#### Functional MRI

fMRI data were preprocessed with FSL 5 (RRID:SCR_002823)^94,95^. Pre-processing included motion-correction with spatial smoothing (7 mm full-width at half maximum, Gaussian kernel) and bandpass filtering (.01-.10 Hz). We registered functional images to participant-specific T1 images, and from T1 to 2mm standard space (MNI 152_T1) using FLIRT. We then masked the functional data with the GM tissue prior provided in FSL (thresholded at probability > 0.37). We detrended the data (up to a cubic trend) using the SPM_detrend function in SPM8. We also utilized extended pre-processing steps to further reduce data artifacts^5,96,97^. Specifically, we subsequently examined all functional volumes for artifacts via independent component analysis (ICA) within-run, within-person, as implemented in FSL/MELODIC^98^. Noise components were identified according to several key criteria: a) Spiking (components dominated by abrupt time series spikes); b) Motion (prominent edge or “ringing” effects, sometimes [but not always] accompanied by large time series spikes); c) Susceptibility and flow artifacts (prominent air-tissue boundary or sinus activation; typically represents cardio/respiratory effects); d) White matter (WM) and ventricle activation^99^; e) Low-frequency signal drift^100^; f) High power in high-frequency ranges unlikely to represent neural activity (≥ 75% of total spectral power present above .10 Hz;); and g) Spatial distribution (“spotty” or “speckled” spatial pattern that appears scattered randomly across ≥ 25% of the brain, with few if any clusters with ≥ 80 contiguous voxels [at 2×2×2 mm voxel size]). Examples of these various components we typically deem to be noise can be found in the supplementary material of Garrett et al^6^. By default, we utilized a conservative set of rejection criteria; if manual classification decisions were challenging due to mixing of “signal” and “noise” in a single component, we generally elected to keep such components. Three independent raters of noise components were utilized; > 90% inter-rater reliability was required on separate data before denoising decisions were made on the current data. To enable semi-automated data denoising using FSL FIX^101,102^, we manually classified 30% of participant data to provide a noise component training set. Features from the noise component training set were then extracted and used to detect noise components from the remaining 70% of participant data via FIX. Upon evaluating the automated labelling for several subjects against our manual decisions, we used a FIX threshold of 60, which permitted a best match to manual decisions of two independent raters. Components identified as artifacts were then regressed from corresponding fMRI runs using the *regfilt* command in FSL. We found previously that these additional preprocessing steps had dramatic effects on the predictive power of SD_BOLD_ in past research, effectively removing 50% of the variance still present after traditional preprocessing steps, while simultaneously doubling the predictive power of SD_BOL_^96^. Critically, our recent work also suggests that when such denoising approaches are applied, age differences in SD_BOLD_ remain robust to multiple vascular controls measured via dual-echo ASL-BOLD using carbogen-based hypercapni^103^. The present sample only contains a narrow age range of older adults (64-68 years), further minimizing the potential impact of aging-based differences in vasculature that may be more present in samples with wider age ranges.

### Network parcellations

To provide full coverage over striatal and thalamic regions, we first created a joint MNI152 registered mask of bilateral striatal (from the Basal Ganglia Human Area Template (BGHAT) atlas^104^) and thalamic regions (from the thalamic nucleus-specific Morel atlas^41^). To examine the relative importance of the striato-thalamic system vs. cortical networks, we also utilized the Yeo 7^105^ parcellation to estimate visual, fronto-parietal (FPN), dorsal attention (DAN), and default-mode networks (DMN). In particular, the FPN is classically assumed to be crucial for parametric working-memory tasks^64–67^, providing perhaps the most flexible hub in the brain specifically for cognitive control and working memory storage^66,106^.

### Brain measures

#### *Voxel-wise estimates of SD*_*BOLD*_

To calculate SD_BOLD_, we first performed a block normalization procedure to account for residual low frequency artifacts. We normalized all blocks for each condition such that the overall 4D mean across brain and block was 100. For each voxel, we then subtracted the block mean and concatenated across all blocks. Finally, we calculated voxel standard deviations across this concatenated time series (Garrett *et al*. 2010). All models described below were run on grey matter (GM) only, after a standard GM mask derived from the MNI152 average brain was applied to each 4D image set.

#### “Temporal” PCA dimensionality as an estimate of functional integration

Building on our previous use of PCA dimensionality to identify spatially coherent networks ^20^, in the current paper, we utilized “temporal” principal components analysis (PCA) as our primary within-subject network dimensionality estimation. Here, separately for each n-back condition and network (i.e., striato-thalamic, visual, FPN, DAN, and DMN), a correlation matrix was estimated for all timepoint pairs (across voxels) from each within-subject data matrix. This correlation matrix was then decomposed using PCA,

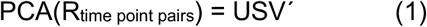

where U and V are the left and right eigenvectors, and S is a diagonal matrix of eigenvalues. We then counted the number of dimensions it took to capture 90% of the within-subject data. Because the S matrix represents the eigenvalues of the solution, and each eigenvalue is proportional to the variance accounted for in the entire decomposition, we summed eigenvalues until 90% of the total variance was reached. In effect, the fewer dimensions it takes to capture 90% of a given subject’s data, the fewer distinct (orthogonal) temporal periods (or temporal “modes”) there are in their time series, suggesting a unification/integration of temporal processing during a given n-back condition. For a comprehensive and direct comparison of information captured by spatial and temporal approaches to component estimation in fMRI, see Smith et al.^107^.

### Statistical modeling

#### Handling of outliers

Prior to model runs, we ran each variable of interest through a modified Winsorization^108^ procedure; all values beyond +-2.5SDs from the mean were flagged as outliers and “Winsorized” by assigning each outlier the closest non-outlying value. All variables were then Z-transformed to minimize collinearity issues that could arise when simultaneously estimating main effects and interactions within the regression models reported for Target 3. All regressions in Target 3 were also run with and without removal of multivariate outliers (see Table S6). We estimated outliers using Cook’s distance; values > 4/n (i.e., 4/152 = .026) were initially flagged as outliers, but only a maximum of 5% of the total sample (= 7 cases) was dropped from any model run.

#### Plotting of interactions

All estimated two-way interactions were between continuous variables (see Table S3). To depict the visual form of such interactions (see Figures 6 and 7), we used a typical point estimate approach^109,110^. Because of the continuous nature of the variables forming these interaction terms, point estimate error bars are, by definition, not estimable. Instead, the unique effect size (pr = partial correlation) of the interaction term estimated within each regression model serves as the relevant marker of “error.”

#### Partial Least Squares

To examine multivariate relations between working memory-based SD_BOLD_ modulation and DA, we utilized behavioral PLS analysis (Mcintosh et al. 1996; Krishnan et al. 2011). This modelling form begins by calculating a between-subject correlation matrix (CORR) between (1) each voxel’s SD_BOLD_ modulation value (i.e., 3-back SD_BOLD_ minus 2-back SD_BOLD_) and (2) D_2_ binding potential (see Results). CORR is then decomposed using singular value decomposition (SVD).

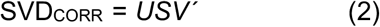

This decomposition produces a left singular vector of offline task weights (*U*), a right singular vector of brain voxel weights (*V*), and a diagonal matrix of singular values (*S*). A single estimable latent variable (LV) results that represents the relations between SD_BOLD_ modulation and DA. This LV contains a spatial activity pattern depicting the brain regions that show the strongest relation to DA identified by the LV. Each voxel weight (in *V*) is proportional to the voxel-wise correlation between DA and SD_BOLD_ modulation.

Significance of detected relations was assessed using 1000 permutation tests of the singular value corresponding to the LV. A subsequent bootstrapping procedure revealed the robustness of within-LV voxel saliences across 1000 bootstrapped resamples of the data (Efron and Tibshirani 1993). By dividing each voxel’s weight (from *V*) by its bootstrapped standard error, we obtained “bootstrap ratios” (BSRs) as non-parametric, normalized estimates of robustness. For the whole brain analysis, we thresholded BSRs at values of ±3.00 (which exceeds a 99% confidence interval).

We also obtained a summary measure of each participant’s robust expression of a particular LV’s spatial pattern (a within-person “brain score”) by multiplying the model-based vector of voxel weights (*V)* by each subject’s vector of voxel SD_BOLD_ upregulation values (*Q*), producing a single within-subject value,

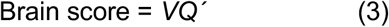

### Code sharing

All code, masks/parcellations, and (un)thresholded brain maps will be available on Github upon final publication at: https://github.com/LNDG/Garrett_etal_2022_COBRA.

