## Supplementary material for "Dynamic regulation of neural variability during working memory reflects dopamine, functional integration, and decision-making": Supp Material

**SUPPLEMENTAL MATERIAL**

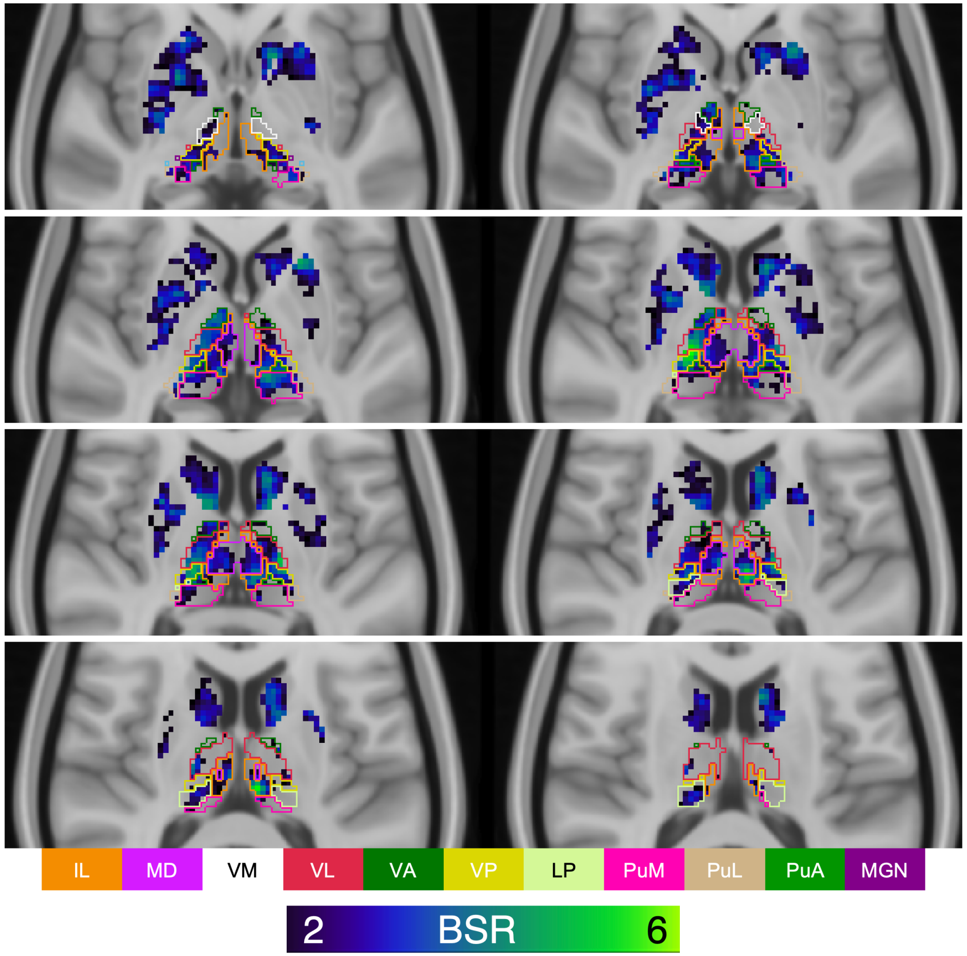


*Figure S1: Axial slice view of the PLS model result depicted in Figure 2 (main paper) with Morel thalamic atlas overlaid.* Blue-green voxels are those representing correlation between working memory load-based $SD_{BOLD}$ modulation and $D_2$ binding. IL = intralaminar. MD = mediodorsal. VM = ventromedial. VL = ventrolateral. VA = ventroanterior. VP = ventroposterior. LP = lateralposterior. PuM = pulvinar, medial. PuL = pulvinar, lateral. MGN = medial geniculate nucleus. BSR = bootstrap ratio.

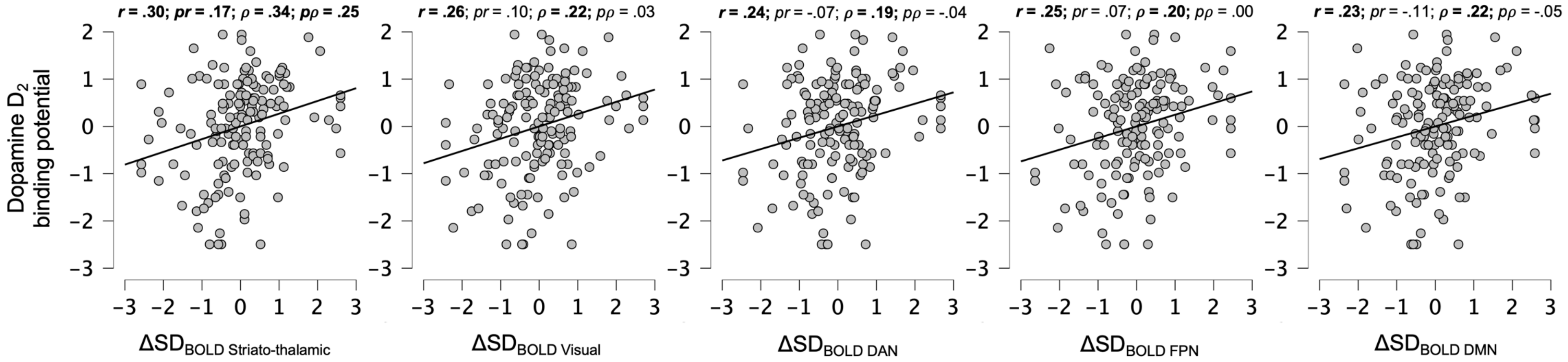


*Figure S2: Correlations between $SD_{BOLD}$ modulation and dopamine $D_2$ binding potential in each a priori network.* Although a positive association between $SD_{BOLD}$ modulation and dopamine was present in every network, partial correlation analyses revealed that this association was uniquely robust in the striato-thalamic system. *pr* = partial Pearson's correlation. *pρ* = partial Spearman's Rho. Statistically significant effects are noted in bold (all statistics are summarized in Table S1).

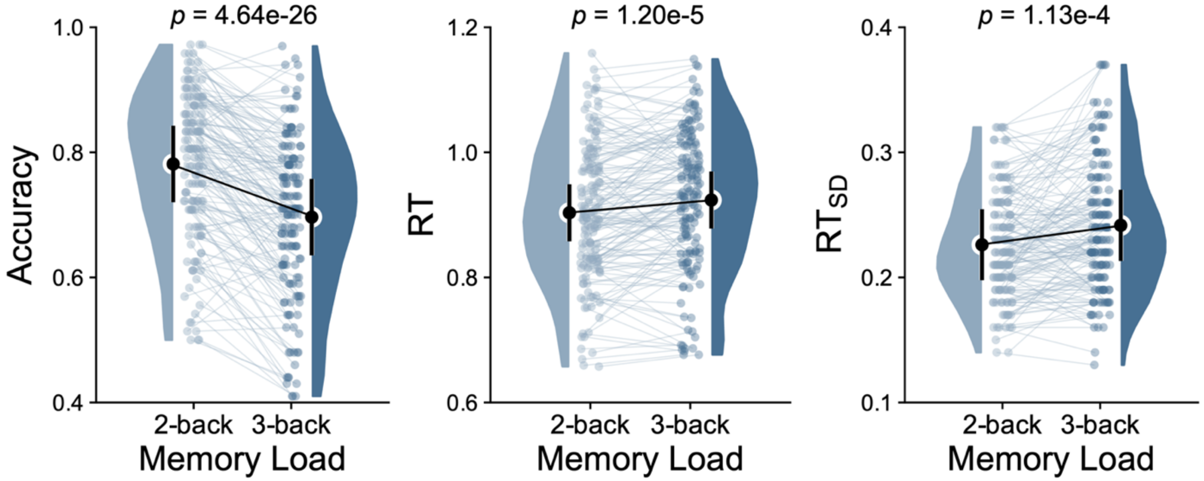


*Figure S3: Accuracy and reaction time (RT) estimates for 2- and 3-back.*

#### 2-back, target absent

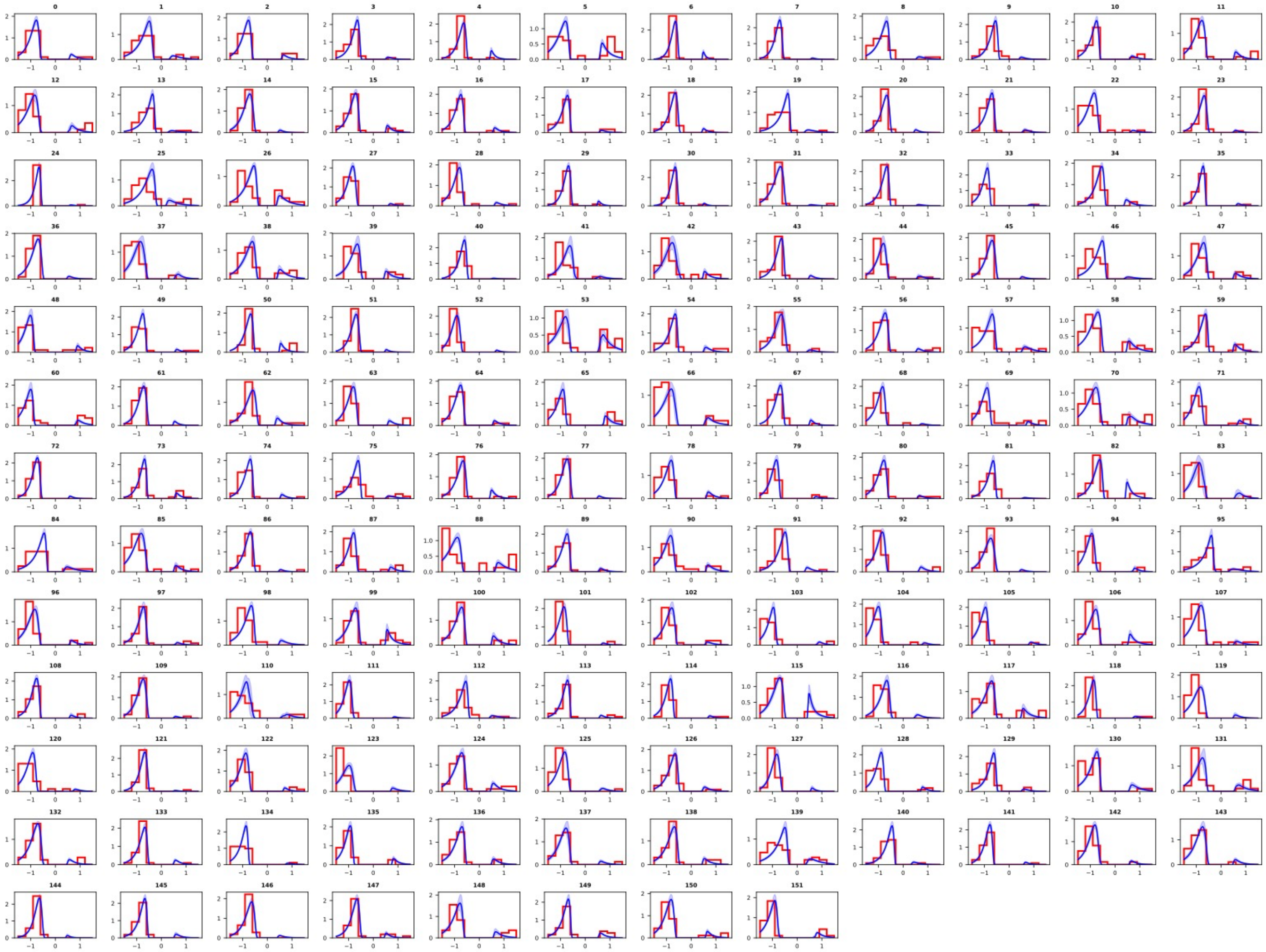

#### 2-back, target present

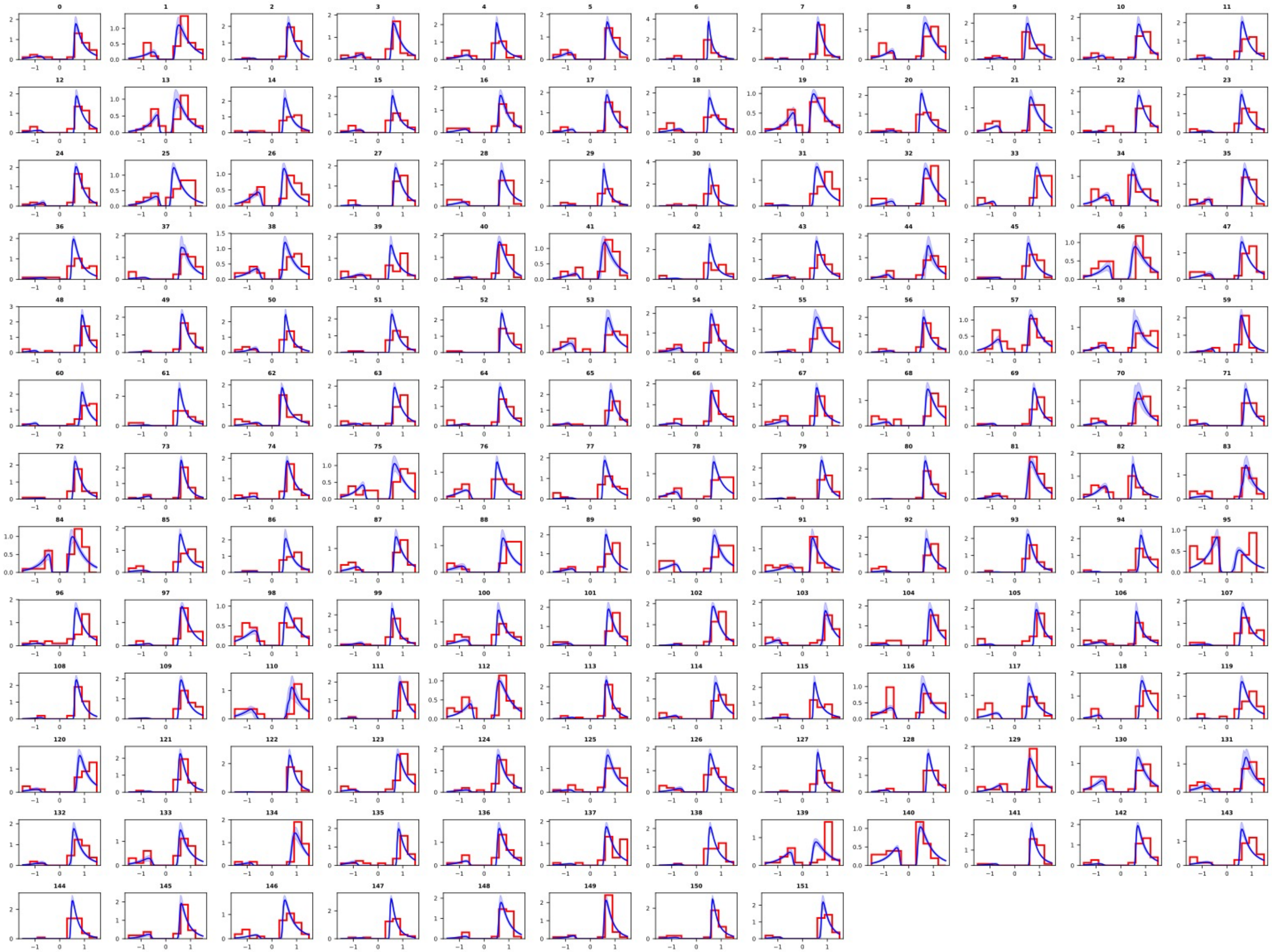

#### 3-back, target absent

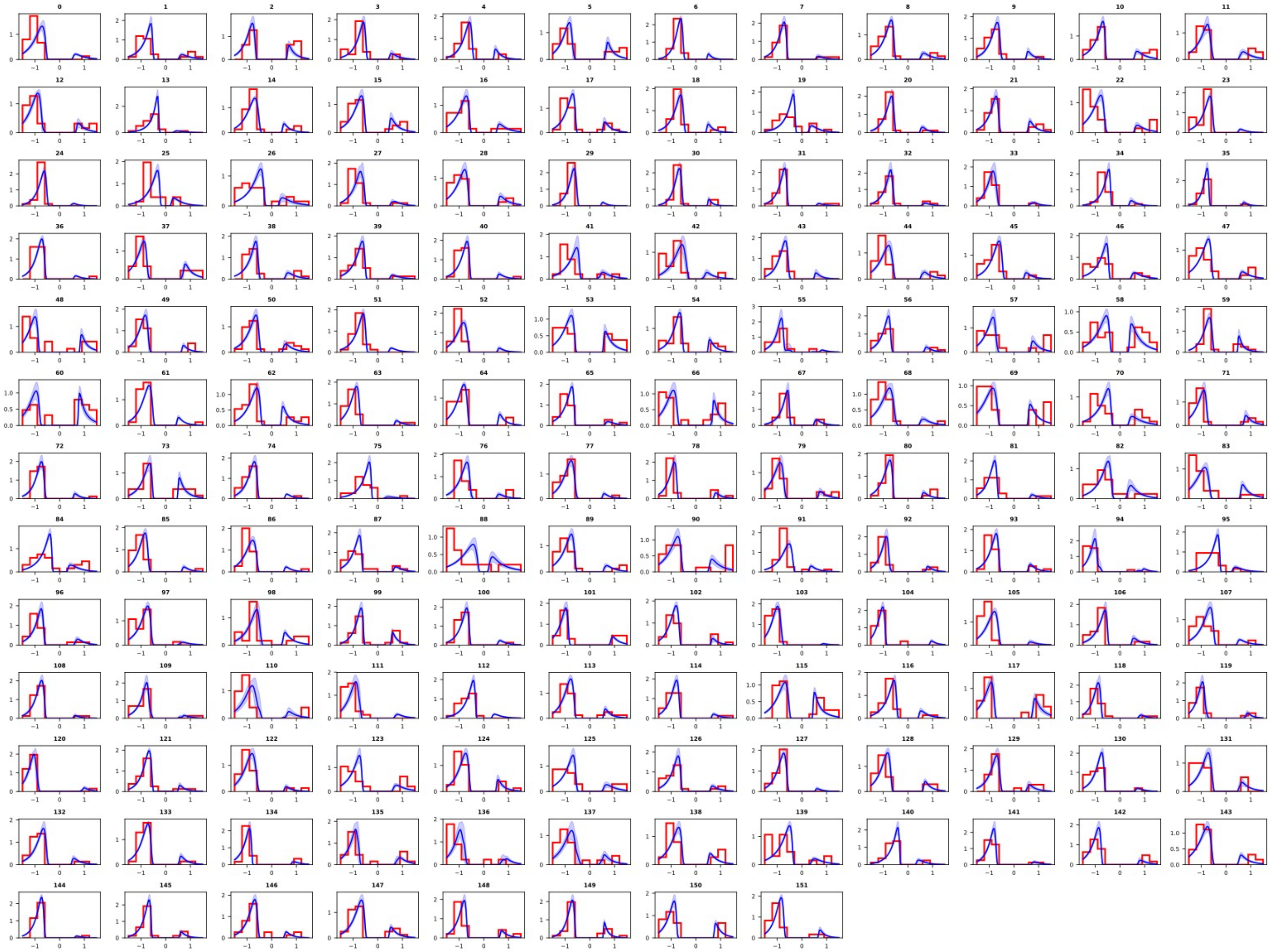

### 3-back, target present

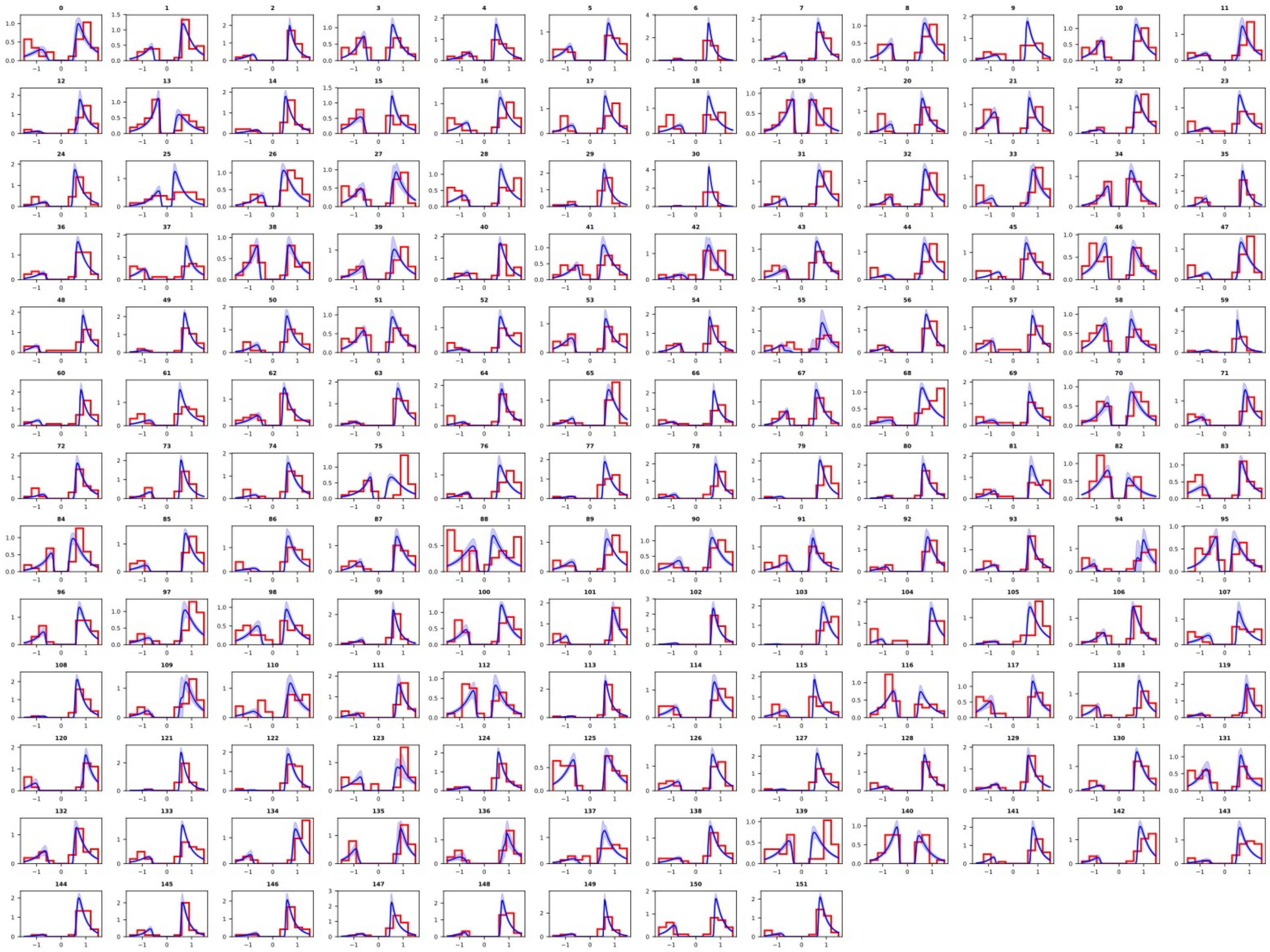

*Figure S4: Reaction time (RT) histograms (red lines) for each subject (N = 152 subjects) with modeling fit (drift rate, boundary separation, non-decision time, starting point bias, drift bias) overlaid (blue lines, error bars are 1 SD) for the 2-back and 3-back conditions.* HDDM "stimulus coding" was used, which fits the DDM separately for target absent (no match) trials and target present (match) trials to enable estimation of decision bias parameters. Fits are shown for both trial types for both conditions. "No" and “Yes” RTs and model fits are negatively and positively signed, respectively. Horizontal axis, RT in seconds; vertical axis, probability density.

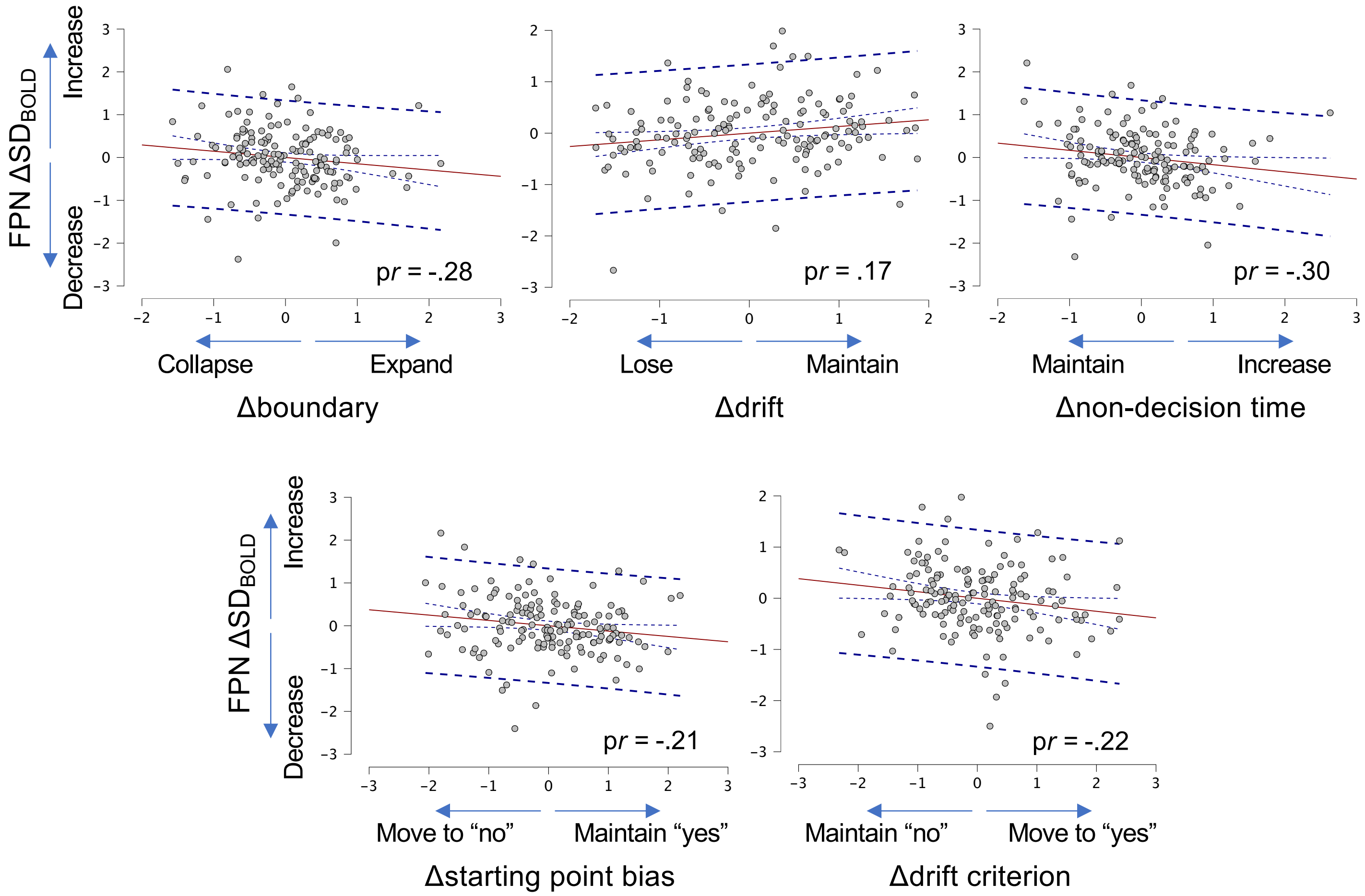


*Figure S4: Main effects of DDM parameters on $SD_{BOLD}$ modulation in the fronto-parietal network (FPN).*

*Table S1: Descriptive statistics and repeated-measures ANOVA (RMANOVA) results for all behavioral parameters.* SDT = signal detection-theoretic. a = boundary; v = drift rate; t = non-decision time; z = starting point bias; dc = drift criterion (drift bias).

| | | | | | | | | RMANOVA models | | |
|---|---|---|---|---|---|---|---|---|---|---|
| | Variable | Load | Mean | Median | SD | Min | Max | *F* | *p* | Partial $\eta^2$ |
| Model-free estimates | Accuracy | 2-back | 0.87 | 0.88 | 0.08 | 0.61 | 0.99 | - | - | - |
| | | 3-back | 0.79 | 0.80 | 0.10 | 0.50 | 1.00 | **168.07** | **4.64E-26** | **0.54** |
| | $RT_{mean}$ | 2-back | 0.89 | 0.88 | 0.12 | 0.62 | 1.20 | - | - | - |
| | | 3-back | 0.92 | 0.93 | 0.13 | 0.61 | 1.17 | **20.60** | **1.20E-05** | **0.12** |
| | $RT_{SD}$ | 2-back | 0.24 | 0.24 | 0.05 | 0.14 | 0.38 | - | - | - |
| | | 3-back | 0.25 | 0.24 | 0.06 | 0.13 | 0.43 | **15.75** | **1.13E-04** | **0.10** |
| DDM parameters | a | 2-back | 1.67 | 1.66 | 0.13 | 1.4 | 1.98 | - | - | - |
| | | 3-back | 1.55 | 1.54 | 0.12 | 1.27 | 1.85 | **127.57** | **7.92E-22** | **0.46** |
| | v | 2-back | 1.52 | 1.54 | 0.37 | 0.61 | 2.33 | - | - | - |
| | | 3-back | 1.1 | 1.11 | 0.35 | 0.24 | 1.9 | **335.77** | **3.24E-40** | **0.69** |
| | t | 2-back | 0.49 | 0.47 | 0.11 | 0.2 | 0.75 | - | - | - |
| | | 3-back | 0.49 | 0.49 | 0.14 | 0.18 | 0.83 | 1.23 | 0.27 | 0.01 |
| | z | 2-back | 0.58 | 0.59 | 0.05 | 0.48 | 0.69 | - | - | - |
| | | 3-back | 0.56 | 0.56 | 0.05 | 0.45 | 0.65 | **61.11** | **8.58E-13** | **0.29** |
| | dc | 2-back | -0.43 | -0.43 | 0.13 | -0.74 | -0.12 | - | - | - |
| | | 3-back | -0.38 | -0.37 | 0.13 | -0.66 | -0.12 | **27.97** | **4.25E-07** | **0.16** |

*Table S2: Bivariate correlation table between offline measures (education, the Digit symbol subtest from the Wechsler Adult Intelligence Scale (WAIS), Working Memory (WM) and Speed) and each DDM parameter.*

| | Education | Digit Symbol | WM | Speed |
|---|---|---|---|---|
| 2-back *a* | 0.125 | 0.069 | 0.145 | -0.083 |
| 3-back *a* | 0.031 | 0.11 | 0.076 | -0.095 |
| 2-back *v* | 0.04 | 0.243** | 0.371*** | 0.227** |
| 3-back *v* | -0.096 | 0.198* | 0.305*** | 0.21** |
| 2-back *t* | -0.121 | -0.195* | -0.168* | -0.096 |
| 3-back *t* | -0.122 | -0.154 | -0.096 | -0.079 |
| 2-back *z* | 0.019 | 0.131 | 0.229** | 0.179* |
| 3-back *z* | -0.133 | 0.025 | 0.051 | 0.102 |
| 2-back *dc* | -0.221** | -0.317*** | -0.403*** | -0.181* |
| 3-back *dc* | -0.29*** | -0.18* | -0.288*** | -0.164* |

*Table S3: Network-specific models of $\Delta SD_{BOLD}$*. All deltas represent n-back load 3 minus load 2. All ΔPCAdim estimates are also network-specific, matching the network of the dependent variable in each model.

| | | N=152 | | | | | | N=145 (w/ Cook's outliers removed) | | | | | |
|---|---|---|---|---|---|---|---|---|---|---|---|---|---|
| **Dependent variable: $\Delta SD_{BOLD}$** | **Independent variables** | ***t*** | ***p*** | **Zero-order** | **Partial** | **Part** | **VIF** | ***t*** | ***p*** | **Zero-order** | **Partial** | **Part** | **VIF** |
| Striato-thalamic | Dopamine | 3.35 | **1.04E-03** | 0.30 | 0.27 | 0.18 | 1.09 | 3.91 | **1.49E-04** | 0.34 | 0.32 | 0.20 | 1.11 |
| | Δa | 0.14 | 0.89 | -0.03 | 0.01 | 0.01 | 2.33 | -0.35 | 0.73 | -0.04 | -0.03 | -0.02 | 2.33 |
| | Δv | -0.08 | 0.94 | 0.01 | -0.01 | 0.00 | 1.26 | 0.07 | 0.94 | 0.00 | 0.01 | 0.00 | 1.27 |
| | Δt | 0.25 | 0.80 | -0.02 | 0.02 | 0.01 | 2.21 | -0.37 | 0.71 | -0.04 | -0.03 | -0.02 | 2.21 |
| | Δz | -1.57 | 0.12 | -0.15 | -0.13 | -0.08 | 1.21 | -1.01 | 0.32 | -0.11 | -0.09 | -0.05 | 1.17 |
| | Δdc | -2.51 | **0.01** | -0.03 | -0.21 | -0.13 | 1.17 | -3.33 | **1.11E-03** | -0.12 | -0.28 | -0.17 | 1.14 |
| | ΔPCAdim | -12.08 | **1.86E-23** | -0.71 | -0.71 | -0.64 | 1.10 | -12.64 | **1.54E-24** | -0.72 | -0.74 | -0.64 | 1.09 |
| | Dopamine * Δdc | -2.56 | **0.01** | -0.17 | -0.21 | -0.13 | 1.05 | -2.50 | **0.01** | -0.15 | -0.21 | -0.13 | 1.05 |
| | ΔPCAdim * Δa | 2.49 | **0.01** | 0.08 | 0.21 | 0.13 | 1.97 | 2.86 | **4.92E-03** | 0.12 | 0.24 | 0.15 | 1.92 |
| | ΔPCAdim * Δv | -2.14 | **0.03** | -0.10 | -0.18 | -0.11 | 1.27 | -2.21 | **0.03** | -0.06 | -0.19 | -0.11 | 1.28 |
| | ΔPCAdim * Δt | 1.85 | 0.07 | 0.08 | 0.15 | 0.10 | 2.10 | 2.46 | **0.02** | 0.08 | 0.21 | 0.13 | 1.98 |
| Yeo1_Visual | Dopamine | 3.20 | **1.66E-03** | 0.26 | 0.26 | 0.18 | 1.03 | 3.36 | **1.02E-03** | 0.28 | 0.28 | 0.19 | 1.03 |
| | Δa | -1.35 | 0.18 | -0.08 | -0.11 | -0.08 | 2.19 | -1.66 | 0.10 | -0.02 | -0.14 | -0.09 | 2.16 |
| | Δv | 1.08 | 0.28 | 0.03 | 0.09 | 0.06 | 1.21 | 1.23 | 0.22 | 0.05 | 0.10 | 0.07 | 1.18 |
| | Δt | -0.58 | 0.56 | 0.06 | -0.05 | -0.03 | 2.03 | -1.21 | 0.23 | 0.00 | -0.10 | -0.07 | 2.01 |
| | Δz | -0.87 | 0.39 | -0.12 | -0.07 | -0.05 | 1.21 | -0.99 | 0.32 | -0.12 | -0.08 | -0.05 | 1.19 |
| | Δdc | -1.49 | 0.14 | -0.07 | -0.12 | -0.09 | 1.14 | -1.84 | 0.07 | -0.08 | -0.16 | -0.10 | 1.15 |
| | ΔPCAdim | -11.39 | **7.74E-22** | -0.69 | -0.69 | -0.65 | 1.04 | -12.20 | **1.23E-23** | -0.72 | -0.72 | -0.68 | 1.07 |
| Yeo3_DAN | Dopamine | 3.31 | **1.18E-03** | 0.24 | 0.27 | 0.17 | 1.04 | 3.63 | **3.95E-04** | 0.23 | 0.30 | 0.17 | 1.04 |
| | Δa | -2.59 | **0.01** | -0.11 | -0.21 | -0.13 | 2.20 | -2.87 | **4.80E-03** | -0.12 | -0.24 | -0.14 | 2.11 |

| | | | | | | | | | | | | | |
|---|---|---|---|---|---|---|---|---|---|---|---|---|---|
| | Δv | 1.93 | 0.06 | 0.05 | 0.16 | 0.10 | 1.24 | 2.31 | **0.02** | 0.04 | 0.19 | 0.11 | 1.20 |
| | Δt | -1.63 | 0.11 | 0.05 | -0.13 | -0.08 | 2.05 | -1.67 | 0.10 | 0.09 | -0.14 | -0.08 | 2.01 |
| | Δz | -0.50 | 0.62 | -0.14 | -0.04 | -0.03 | 1.22 | 0.20 | 0.84 | -0.09 | 0.02 | 0.01 | 1.18 |
| | Δdc | -3.10 | **2.35E-03** | -0.10 | -0.25 | -0.16 | 1.14 | -3.42 | **8.21E-04** | -0.13 | -0.28 | -0.16 | 1.13 |
| | ΔPCAdim | -12.98 | **6.12E-26** | -0.72 | -0.74 | -0.67 | 1.10 | -14.91 | **2.09E-30** | -0.75 | -0.79 | -0.72 | 1.08 |
| | Dopamine * Δa | -2.26 | **0.03** | -0.23 | -0.19 | -0.12 | 1.06 | -3.37 | **9.73E-04** | -0.24 | -0.28 | -0.16 | 1.05 |
| Yeo6_FPN | Dopamine | 2.90 | **4.27E-03** | 0.25 | 0.24 | 0.16 | 1.04 | 3.26 | **1.42E-03** | 0.26 | 0.27 | 0.16 | 1.04 |
| | Δa | -1.82 | 0.07 | -0.06 | -0.15 | -0.10 | 2.23 | -3.35 | **1.04E-03** | -0.11 | -0.28 | -0.16 | 2.43 |
| | Δv | 2.17 | **0.03** | 0.03 | 0.18 | 0.12 | 1.22 | 2.05 | **0.04** | -0.02 | 0.17 | 0.10 | 1.20 |
| | Δt | -2.14 | **0.03** | 0.02 | -0.18 | -0.12 | 2.09 | -3.71 | **2.98E-04** | 0.02 | -0.30 | -0.18 | 2.26 |
| | Δz | -2.09 | **0.04** | -0.17 | -0.17 | -0.11 | 1.21 | -2.51 | **0.01** | -0.17 | -0.21 | -0.12 | 1.19 |
| | Δdc | -2.18 | **0.03** | -0.08 | -0.18 | -0.12 | 1.17 | -2.65 | **0.01** | -0.08 | -0.22 | -0.13 | 1.19 |
| | ΔPCAdim | -12.54 | **8.44E-25** | -0.70 | -0.72 | -0.68 | 1.07 | -14.63 | **1.05E-29** | -0.74 | -0.78 | -0.72 | 1.07 |
| | ΔPCAdim * Δv | -2.48 | **0.01** | -0.07 | -0.20 | -0.13 | 1.06 | -3.29 | **1.28E-03** | -0.12 | -0.27 | -0.16 | 1.07 |
| Yeo7_DMN | Dopamine | 3.07 | **2.53E-03** | 0.23 | 0.25 | 0.16 | 1.03 | 3.07 | **2.60E-03** | 0.21 | 0.25 | 0.15 | 1.03 |
| | Δa | -1.26 | 0.21 | -0.01 | -0.10 | -0.07 | 2.23 | -0.94 | 0.35 | -0.01 | -0.08 | -0.05 | 2.23 |
| | Δv | 0.93 | 0.35 | 0.02 | 0.08 | 0.05 | 1.21 | 0.13 | 0.90 | -0.02 | 0.01 | 0.01 | 1.20 |
| | Δt | -1.51 | 0.13 | 0.00 | -0.13 | -0.08 | 2.07 | -1.23 | 0.22 | 0.05 | -0.10 | -0.06 | 2.05 |
| | Δz | -0.65 | 0.52 | -0.13 | -0.05 | -0.03 | 1.22 | -0.84 | 0.40 | -0.14 | -0.07 | -0.04 | 1.24 |
| | Δdc | -1.94 | 0.05 | -0.07 | -0.16 | -0.10 | 1.14 | -1.89 | 0.06 | -0.05 | -0.16 | -0.09 | 1.16 |
| | ΔPCAdim | -13.20 | **1.47E-26** | -0.73 | -0.74 | -0.71 | 1.07 | -14.90 | **1.92E-30** | -0.78 | -0.79 | -0.75 | 1.09 |
